# Uncoupling the Effects of Highland Maize Chromosomal Inversion *Inv4m* from Leaf Phosphorus Deficiency Responses

**DOI:** 10.64898/2026.05.22.727253

**Authors:** Fausto Rodríguez-Zapata, Ruthie Locklear, Allison C. Barnes, Nirwan Tandukar, Sergio Pérez-Limón, Melanie Perryman, Miguel A. Piñeros, Jonathan Odilón Ojeda-Rivera, Daniel Runcie, Ruairidh Sawers, Rubén Rellán-Álvarez

## Abstract

Local adaptation of a species involves the selection of adaptive alleles that confer a fitness advantage in their local environment. Inversions prevent recombination between the standard and inverted heterozygous hybrids. Inversions can play a crucial role in local adaptation by locking together a set of co-adapted alleles, acting as supergenes. *Inv4m* is a 13 Mb inversion in maize prevalent in highland maize and highland wild relatives from México. Maize from the highlands of the Trans-Mexican volcanic belt has been shown to be well-adapted to volcanic, acidic soils with low phosphorus availability. *Inv4m* carries several genes involved in P acquisition and utilization. We therefore tested the hypothesis that *Inv4m* contributes to maize adaptation to these environments through enhanced phosphorus acquisition or utilization. Alternatively, *Inv4m* possible adaptive value may operate through constitutive developmental effects independent of nutrient stress responses. To test this hypothesis, we introgressed a highland maize variety from the highlands of Michoacán, México, carrying *Inv4m* into the temperate line B73 and developed Near-Introgression Lines (NILs) carrying *Inv4m*. We then grew NILs carrying the inversion and controls without it in soils with different phosphorus levels and evaluated the fitness effects of the inversion, as well as changes in gene expression using RNA-Seq. Our results show that P starvation elicits highly conserved transcriptomic, lipidomic, and ionomic responses, independently of the *Inv4m* inversion genotype. Therefore, phosphorus deficiency does not seem to be driving the adaptive value of *Inv4m*. Additionally, we observed a phosphorus modulated transcriptional gradient from the collar leaf downward, characterized by a decrease in the expression of photosynthesis genes and an increase in the expression of senescence-associated genes, corresponding to the positional onset and initial stages of sequential leaf senescence. Although the magnitude of the phosphorus response increased with leaf age, we did not observe significant interactions with *Inv4m*. Our multi-omics analysis of the maize phosphorus starvation response identified and characterized two coordinately regulated molecular programs, light harvesting shutdown and accelerated senescence, whose deployment depends on leaf developmental stage, with older leaves below the collar integrating nutrient limitation into the natural progression toward senescence.

## Introduction

MAize was originally domesticated in the tropical lowlands of Mexico. Before its expansion into temperate regions, maize was introduced to the Mexican highlands, where sympatry with highland teosinte *Zea mays ssp. mexicana* (shorthand *mexicana*) likely facilitated the introgression of adaptive alleles from *mexicana*. Teosinte *mexicana* introgression probably facilitated adaptation to temperate zones and further expansion worldwide (Yang *et al*. 2023; Guo *et al*. 2018; Barnes *et al*. 2022).

However, while the average amount of *mexicana* introgression in modern maize is around 18% (Yang *et al*. 2023), not all highland loci are present in temperate maize. Highland-associated chromosomal inversions, such as *Inv4m* and *Inv9f*, are prevalent in highland teosinte populations (Pyhäjärvi *et al*. 2013) and traditional Mexican maize varieties (TVs) (Crow *et al*. 2020; Gonzalez-Segovia *et al*. 2019) but are rare in modern temperate maize.

Chromosomal inversions can contribute to local adaptation by preserving locally adapted alleles across multiple loci and reducing recombination within the inversion (Kirkpatrick and Barton 2006). Genotyping of teosinte populations revealed that *Inv4m* spans 13 Mb and is predominantly found in *mexicana* populations (Pyhäjärvi *et al*. 2013). In Mexican TVs, *Inv4m* genotype is associated with elevation and flowering time (Romero Navarro *et al*. 2017). Additionally, *Inv4m* shows reduced genetic diversity, a clinal relationship with elevation, and is nearly fixed in locations above 2500 m.a.s.l. (Crow *et al*. 2020). The inversion shows suppressed recombination, as confirmed in a biparental cross (Gonzalez-Segovia *et al*. 2019). *Inv4m* demonstrates classic patterns of gene-by-environment interactions indicative of local adaptation. Plants carrying the *Inv4m*-highland allele show delayed flowering at low elevations and earlier flowering at high elevations (Gates *et al*. 2019; Barnes *et al*. 2022).

Despite strong evidence linking *Inv4m* to local adaptation, the physiological processes and environmental factors underlying its adaptive role remain unclear. Furthermore, the specific genes within *Inv4m* that drive local adaptation are largely unidentified. Previous research has shown that *Inv4m*-highland upregulates photosynthesis genes in response to cold at the seedling stage (Crow *et al*. 2020) and is associated with earlier flowering in the Mexican highlands, which likely enhances fitness in environments with limited growth-degree accumulation throughout the year (Romero Navarro *et al*. 2017). However, cold is not the only limiting factor for plant growth in the Mexican highlands. Volcanic soils (Andosols), which dominate the Mexican highlands, present an additional constraint. Approximately 95% of natural Andosol profiles in Mexico are found above 2000 m.a.s.l. (Paz-Pellat and Velázquez-Rodríguez 2018; INEGI 2013). These soils are characterized by high phosphorus retention (Krasilnikov *et al*. 2013), which leads to low phosphorus availability for plant uptake (Galván-Tejada *et al*. 2014). MICH21, one of the Mexican highland maize accessions analyzed by (Crow *et al*. 2020), originates from the Purépecha Plateau, where Andosols and phosphorus-efficient TVs are common (Paz-Pellat and Velázquez-Rodríguez 2018; Galván-Tejada *et al*. 2014; Bayuelo-Jiménez *et al*. 2011; Bayuelo-Jiménez and Ochoa-Cadavid 2014). *Inv4m* may contribute to adaptation in the highlands by carrying alleles that enhance the phosphorus starvation response (PSR). For example, the phosphate transporter gene *ZmPho1;2a*, located within *Inv4m*, is a strong candidate for adaptation to low phosphorus availability (Salazar-Vidal *et al*. 2016; Ma *et al*. 2021).

The developmental differentiation of the canopy serves as a framework for understanding the effects of phosphorus stress on the leaves. Annual plants like *Arabidopsis* (Hensel *et al*. 1993) and rice (Mondal and Choudhuri 1984) show sequential or acropetalous senescence (Leopold 1961), where aging progresses from the basal to the apical leaves. It is a common observation that basal maize leaves senesce and die early in development (Dudley and Poethig 1991), establishing the oldest portion of the vertical senescence axis. However, rather than a unidirectional gradient, the vertical profile of physiological activity in maize follows a bell-shaped curve, whose peak depends on the developmental stage (Ciganda *et al*. 2008). After flowering, chlorophyll content (Ciganda *et al*. 2008), leaf area and longevity (Valentinuz and Tollenaar 2004), nitrogen (Wei *et al*. 2025), and water content (Gao *et al*. 2023) first increase and then decline with leaf position, peaking near the ear leaf. During the vegetative phase, peak chlorophyll content and metabolic activity occur in the uppermost fully expanded ‘collar’ leaf (Ciganda *et al*. 2008). While the steepness of this gradient is genotype dependent (Gao *et al*. 2023), this architecture generally drives an ‘outside-in’ senescence pattern where aging proceeds from both the top and bottom of the canopy toward the middle (Wei *et al*. 2025) of the plant close to the main ear. This physiological architecture ensures that leaves closest to the ear, the primary sink in the reproductive phase, retain longevity and photosynthetic activity the longest (Valentinuz and Tollenaar 2004). The delay in senescence near the ear aligns with the principle of proximity allocation, which minimizes transport distances for assimilates and optimizes energy use during the critical grain-filling period (Wei *et al*. 2025; Valentinuz and Tollenaar 2004). In contrast, the collar leaf, the youngest and fully developed leaf during vegetative growth, is a source of photosynthates rather than a sink (Evert *et al*. 1996). This pattern might be indicative of its proximity to the main shoot carbon sinks of the vegetative phase: the fastest elongating internodes and immature leaves (Zhao *et al*. 2022).

Disentangling the phosphorus starvation response from this autonomous sequential senescence gradient is complicated by the known interaction between phosphorus limitation and the genetic circuitry of senescence. In *Arabidopsis*, the genetic regulation of phosphorus starvation response and leaf senescence are intertwined, with unambiguous phenotypic consequences following genetic perturbation. The transcription factor PHR1 directly binds to and activates senescence-associated genes, including *RNS1, PAP17*, and *PLDζ2*, with overexpression accelerating leaf senescence and facilitating phosphorus transfer to young tissues (Zhang *et al*. 2024). Phospholipases PLD*ζ*2 and NPC4 are highly induced during senescence, and their knockouts delay senescence while impairing phosphorus remobilization (Yang *et al*. 2024). Similarly, knocking out the purple acid phosphatase AtPAP26 delays leaf senescence, impairs phosphorus remobilization efficiency and reduces seed phosphorus concentrations (Robinson *et al*. 2012). These studies establish a conserved mechanism wherein phosphorus recycling through membrane phospholipid hydrolysis and enzymatic phosphorus scavenging directly promote senescence. In maize, phosphorus starvation studies detect molecular machinery homologous to that of *Arabidopsis*: ribonucleases, phosphatases, and membrane lipid remodeling enzymes (Zhang *et al*. 2014b; Torres-Rodríguez *et al*. 2024) are upregulated and proteomic analyses show increased antioxidant enzymes and altered photosynthetic proteins (He *et al*. 2022; Zhang *et al*. 2014a). However, details beyond those have yet to be revealed in maize. There is a lack of mechanistic knowledge and inconsistent physiological evidence regarding the effect of P deficiency on sequential senescence. Field studies in maize report variable outcomes, ranging from delayed senescence in lower leaves (Colomb *et al*. 2000), to only slight whole-plant effects (Plénet *et al*. 2000), or negligible differences in stalk senescence (Russo and Pappelis 1995). Evidence from a greenhouse assay suggests that nitrogen remobilization from older to younger leaves occurs in plants experiencing phosphorus limitation (Usuda 1995). Specifically, older leaves show a noticeable reduction in both nitrogen and chlorophyll content; however, the phosphorus treatment had little to no impact on the nitrogen and chlorophyll content of the young leaves.

This suggests that phosphorus restriction regulates leaf senescence differentially, accelerating it in older leaves in order to delay it in younger leaves. These variable results may also reflect differences in sampling strategies and developmental timing.

In this study, we aimed to understand the physiological and molecular effects of *Inv4m* and to identify candidate genes within the inversion that could elucidate its adaptive role. Specifically, we tested whether *Inv4m*-highland contributes to adaptation to low phosphorus availability. To achieve this, we backcrossed MICH21, a Mexican highland TV carrying *Inv4m*, into the B73 genetic background for eight generations, generating Near Isogenic Lines (NILs). These NILs were grown under temperate field conditions with two phosphorus treatments. We sampled specific leaf positions at a defined developmental stage (V13, approximately 10-16 days pre-flowering) during the onset of sequential senescence, likely capturing transient molecular responses that whole-plant measurements or variable-timing sampling protocols might miss.

Our multi-omics analysis indicates that the metabolic response to this stress was consistent between the *Inv4m* NILs and the B73 control at the molecular and organismic level. We did not find a specific interaction between *Inv4m* and P deficiency; instead, the molecular response was driven by the leaf’s position on the plant and nutrient limitation. Older leaves showed enhanced stress signatures distinct from those of younger tissues, highlighting the compounding effect of developmental stage on the starvation response. Ultimately, *Inv4m* did not significantly alter these metabolic pathways, suggesting that its adaptive value in highland environments may stem from its constitutive effects on phenology rather than a specific modification of phosphorus metabolism.

## Results

### Phosphorus starvation has strong maize phenotypic effects independent of Inv4m

Phosphorus deficiency delayed flowering, reduced biomass, and decreased yield (Fig 1) in sibling NILs homozygous for the B73 reference (CTRL) and *Inv4m*-MICH21 karyotypes alike. Under −*P* conditions, anthesis and silking occurred more than three days later relative to +*P* (Fig 1A and B; anthesis: 3.60 ± 0.26 days, FDR = 1.1 × 10^−18^;silking: 3.42 ± 0.23 days, FDR = 8.1 × 10^−20^; marginal effect estimate ± S.E, FDR adjusted across the 144-test phenotype family). In reduced phosphorus, the 50-kernel weight decreased by nearly 18% (−1.86 ± 0.37 g, FDR = 4.1 × 10^−5^, Fig 1C). Under −P, 50-kernel weight decreased in both CTRL (−1.80 g, FDR = 0.005) and *Inv4m* (−1.92 g, FDR = 0.002) plants, total kernel weight per plant remained statistically unchanged in either genotype, and total kernel number increased only in the marginal contrast pooled over genotype (+58.7, FDR = 0.044; S1 Fig). Biomass accumulation was diminished under phosphorus starvation at all measured time points (Fig 1G): stover dry weight declined by 5.09 ± 0.71 g at 40 DAP (FDR = 1.1 × 10^−8^), 16.45 ± 1.11 g at 50 DAP (FDR = 1.1 × 10^−18^), and 27.94 ± 3.47 g at 60 DAP (FDR = 1.4 × 10^−9^). By harvest time (121 DAP), stover biomass remained around 18.5% lower (−18.70 ± 3.27 g, FDR = 2.3 × 10^−6^). Fitted logistic growth curves captured the effect of P-starvation on multiple model parameters(S3 Fig A). −P treatment significantly reduced the area under both the empirical (AUCE; −1.96 ± 0.18 kg × day, *p* = 2.2 × 10^−14^) and logistic growth curves (AUCL; −1.80 ± 0.19 kg × day, *p* = 2.0 × 10^−12^) (S3 Fig A). Additionally, it reduced the fitted maximum stover weight (STW_max_) by approximately 23.00 ± 3.26 g (*p* = 5.6 × 10^−9^) and delayed the time to reach half maximum stover weight (T_1/2_) by 3.49 ± 0.80 days (*p* = 6.0 × 10^−5^), relative to the +P treatment (S3 Fig A). We found no significant difference in the rate of stover biomass accumulation. These phenotypic changes match the canonical maize phosphorus starvation syndrome, indicating a robust physiological response to nutrient limitation. Crucially, no significant genotype-by-phosphorus interactions were detected for the primary agronomic traits, implying that the *Inv4m* inversion did not alter the direction or magnitude of the main phosphorus response (Fig 1).

**Figure 1.**
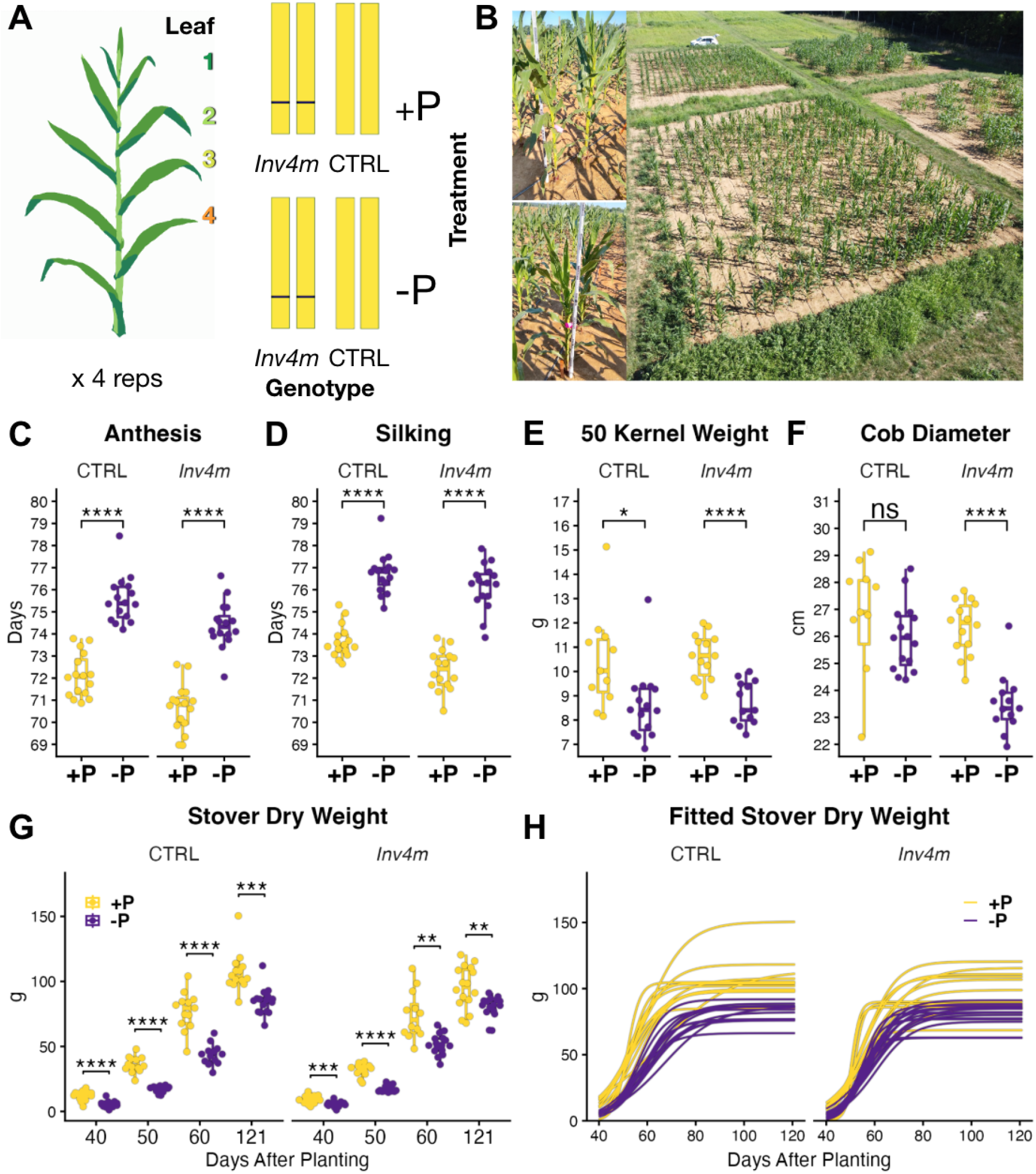
Phosphorus Starvation Response. **(A)** Experimental design schematics showing the four sampled leaves from *Inv4m* and control (CTRL) NILs at 63 DAP (~ V13). An increasing number corresponds to older leaves. **(B)** General appearance of plants at RNA/lipid sampling. *Top* +P, *Botttom* −P treatments. *Right*: Aerial view of the experimental field at Rocksprings, PA. Phosphorus starvation led to delayed anthesis **(C)** and silking **(D)**, reduced 50 kernel weight. Cob diameter **(F)** showed the only significant *Inv4m* genotype dependency, resulting in thinner cobs under phosphorus deficiency. **(G)** Time course of stover dry weight shows lower biomass accumulation under −*P* across all sampling dates for both genotypes. **(H)** Fitted logistic growth curve, each line corresponds to a plot. *FDR* adjusted contrast *t-test* significance: *n*.*s*. not significant, *p <* 0.05 (*), *p <* 0.01 (**), *p <* 0.001 (***), *p <* 0.0001 (****).

Nonetheless, we observe a significant *G* × *E* interaction effect for cob diameter, a secondary reproductive trait. Specifically, while the cob diameter of control lines was decreased less in P-starved plants (−0.19 ± 0.70 cm, FDR = 0.85), the *Inv4m* plants grew a cob 10.7% thinner under nutrient limitation (−2.81 ± 0.68 cm, FDR = 6.2 × 10^−4^; per-genotype effect estimate ± S.E, FDR adjusted across the 144-test phenotype family, Fig 1F). Aside from cob diameter, the effects of *Inv4m* were independent of phosphorus conditions and smaller than those of phosphorus starvation.

Pooled across treatments, *Inv4m* plants flowered earlier (anthesis: −1.31 ± 0.26 days, FDR = 2.5 × 10^−5^; silking: −0.93 ± 0.23 days, FDR = 6.0 × 10^−4^) and grew taller by 6.41 ± 1.05 cm (FDR = 5.0 × 10^−7^); marginal *Inv4m* vs CTRL effects from the 144-test phenotype family, S2 Fig. We found a significant linear time dependence for −P relative reduction in biomass (*p* = 0.027) but the time effect was not significant for the inversion (*p* = 0.90), S3 Fig B). The biomass lag of −P plants relative to the +P decreased as plants matured, from 48% at 40 DAP to 18% at harvest. Overall, while phosphorus starvation consistently resulted in severe reproductive and vegetative penalties, we did not find interactions between *Inv4m* and phosphorus deficiency.

### Plant mineral concentrations show major responses to phosphorus starvation but only minor perturbations from the Inv4m inversion

Phosphorus deficiency (−P) induced strong and consisted shifts in mineral accumulation across both genotypes, indicating that the overall ionomic response is largely shared between the *Inv4m* and control lines (Fig 2 A and B). Phosphorus concentrations declined sharply under −P in both stover (effect estimate ± s.e: −1592 ± 85 ppm, *p* = 1.13 × 10^−25^) and seeds (−672 ± 94 ppm, *p* = 1.68 × 10^−8^), accompanied by a strong increase in the seed/stover P ratio (1.99 ± 0.13, *p* = 1.25 × 10^−19^). Zinc levels increased in stover (6.85 ± 1.12 ppm, *p* = 3.24 × 10^−7^), while Ca rose in seed (18.96 ± 3.43 ppm, *p* = 4.35 × 10^−6^), with corresponding changes in Zn and Ca partitioning ratios (−0.23 ± 0.04, *p* = 3.62 × 10^−6^; and +0.0041 ± 0.00085, *p* = 4.17 × 10^−5^, respectively). Sulfur concentrations also increased under −P in both stover (113 ± 21 ppm, *p* = 4.49 × 10^−6^) and seed (79 ± 30 ppm, *p* = 2.96 × 10^−2^), while Mg decreased modestly in seed (−97 ± 30 ppm, *p* = 7.15 × 10^−3^). We found a genotype-dependent response to phosphorus for stover sulfur where *Inv4m* plants accumulated less sulfur under P deficiency than the control line (*G* × *E* interaction, −93.8 ± 29.2 ppm, *p* = 6.50 × 10^−3^).

**Figure 2.**
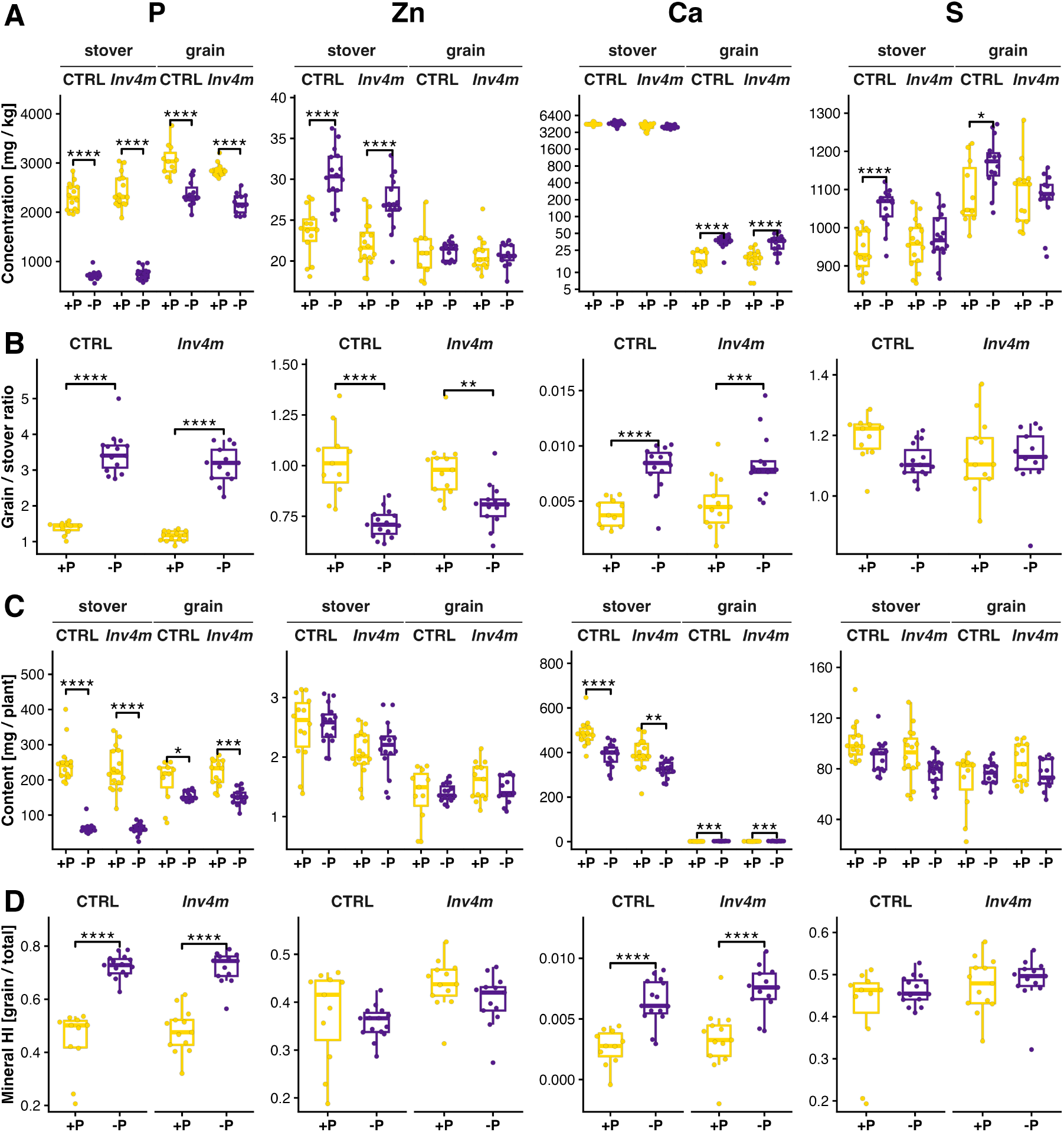
Ionomic responses of *Inv4m* and control (CTRL) maize lines under phosphorus sufficiency (+P) and deficiency (−P). Boxplots for phosphorus (P), zinc (Zn), calcium (Ca), and sulfur (S) show, per genotype, **(A)** element concentrations in stover and grain, **(B)** grain/stover concentration ratios, **(C)** content in stover and grain per plant, and **(D)** mineral harvest index (HI, grain/total content). Ca concentrations are plotted on a log scale. *FDR* adjusted contrast *t-test* significance: *n*.*s*. not significant, *p <* 0.05 (*), *p <* 0.01 (**), *p <* 0.001 (***), *p <* 0.0001 (****). Effect sizes and exact *p* values are reported in S1 Table.

Mineral content per plant and harvest index (Fig 2 C and D) complement the concentration analysis. Phosphorus content decreased in both tissues under −P (stover: −186 ± 16 mg, *p* = 2.0 × 10^−16^; grain: −41 ± 15 mg, *p* = 0.019), and the grain P harvest index rose (+0.28 ± 0.03, *p* = 1.7 × 10^−11^). Calcium content decreased in stover (−97 ± 21 mg, *p* = 4.9 × 10^−5^) and increased in grain (+1.26 ± 0.29 mg, *p* = 1.9 × 10^−4^), with a corresponding rise in Ca harvest index (+0.0037 ± 0.0008, *p* = 4.9 × 10^−5^). Zinc content was statistically unchanged in both tissues (stover: *p* = 0.86; grain: *p* = 0.89). Sulfur content did not change significantly in either tissue after FDR adjustment (stover: −11.4 ± 5.7 mg, *p* = 0.11; grain: +6.2 ± 6.0 mg, *p* = 0.45). For Mg, Mn, K, and Fe (S4 Fig), stover content decreased under −P (Mg: −35 ± 8 mg, *p* = 3.2 × 10^−4^; K: −299 ± 84 mg, *p* = 2.3 × 10^−3^; Mn: −0.66 ± 0.25 mg, *p* = 0.025), while grain content and harvest index were unchanged; Fe showed no significant shifts in either tissue.

When pooled across phosphorus treatments, nine additive *Inv4m* effects on the ionome reached significance (S5 Fig). Calcium in stover was lower in *Inv4m* both as content (−94 ± 21 mg, *p* = 9.0 × 10^−5^) and as concentration (−411 ± 141 ppm, *p* = 0.014). Zinc content in stover decreased (−0.44 ± 0.16 mg, *p* = 0.019), while the Zn harvest index increased (+0.064 ± 0.025, *p* = 0.032). Iron was preferentially allocated to grain: grain content rose (+0.21 ± 0.07 mg, *p* = 0.020), the grain/stover ratio increased (+0.053 ± 0.019, *p* = 0.019), and the Fe harvest index rose (+0.040 ± 0.013, *p* = 0.010). Magnesium grain concentration decreased (−95.6 ± 31.5 ppm, *p* = 0.011), and potassium stover content decreased (−213 ± 84 mg, *p* = 0.034).

### Phosphorus starvation triggers a robust and canonical remodeling of the maize leaf transcriptome

To further understand the effect of *Inv4m*, P-defiency and leaf age on the at the molecular level we performed an RNA-Seq experiment. A multidimensional scaling (MDS) of gene expression (as *log*_2_[CPM], counts per million) captured 38% variance in the first two dimensions (Fig 3A). The first dimension alone explained 26% of variance and is correlated with phosphorus treatment (Pearson *r* = 0.50, *t-test FDR* = 6.15 × 10^−4^). Phosphorus (P) starvation led to a global transcriptional response with a total of 10,606 differentially expressed genes (DEGs, *FDR <* 0.05) out of the 24011 detected in the sampled leaves. The core of the response involved classic mechanisms of P mobilization and reallocation, which are conserved across plant species (Fig 3B, S3 Table).

**Figure 3.**
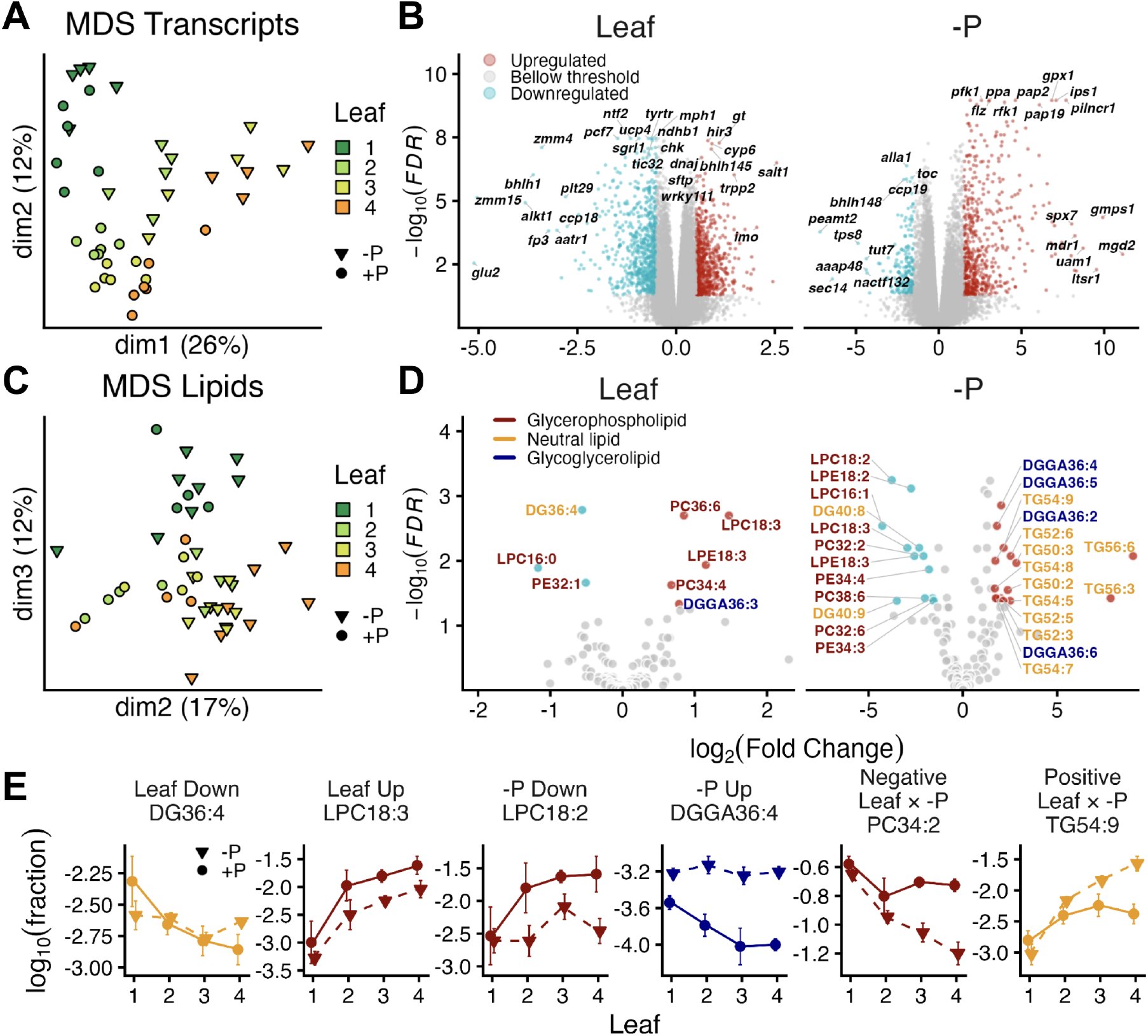
Transcriptomic and lipidomic responses to phosphorus deficiency and leaf developmental stage. (**A**) Multidimensional scaling of transcripts. The MDS plot of log_2_(CPM). (**B**) Volcano Plots of Transcriptomic Main Effects. *Right*: Main transcriptional effect of leaf stage (per-stage increase). A total of 1,431 strong DEGs were identified. The x-axis represents the log_2_(Fold Change) and the y-axis represents the − log_10_(FDR). *Left*: Main transcriptional effect of Phosphorus deficiency (−P treatment). (**C**) Multidimensional scaling of lipids. The MDS plot of log_2_(CPM). (**D**) Volcano plots of lipidomic main effects. *Left*: Main effect of leaf stage on lipids. The axes and thresholds are analogous to those in panel B, highlighting lipids whose abundance is significantly altered by each factor independently of the other. *Right:* Main effect of phosphorus deficiency on lipids. (**E**) Mass Spectrometry signal profiles of the most differentially abundant lipids.

The upregulated genes showed enrichment in cellular response to phosphate starvation (Fisher’s exact test, *FDR* = 9.07 × 10^−11^) (Fig 4A, S5 Table). Top DEGs known to be upregulated under P starvation S2 Table included *Pap19* (*Zm00001eb010130*, − log_10_ *FDR* = 9.7, *log*2 *FC* = 5.99), encoding a purple acid phosphatase that hydrolyzes organic P compounds; *pilncr1* (*Zm00001eb003820*, − log_10_ *FDR* = 9.6, *log*_2_ *FC* = 7.34), a P deficiency-induced long non-coding RNA and precursor to *miR399* (a master regulator of P homeostasis); and *ips1* (*Zm00001eb148590*, − log_10_ *FDR* = 9.3, *log*_2_ *FC* = 7.08) which is a decoy target for *miR399* that prevents it from repressing the *PHO2* transporter, thereby enhancing P uptake efficiency (Du *et al*. 2018). The P-starvation response also involved modification of leaf membrane lipids. Other up-regulated genes included in the overrepresented KEGG set were: several *SPX* family transcription factors, the phosphate transporters *pho1;1* (*Zm00001eb126380*), *pht1* (*Zm00001eb222510*) and *pht7* (*Zm00001eb038730*), which facilitate phosphate up-take and redistribution; and the purple acid phosphatases *Pap1* (*Zm00001eb151650*) and *Pap14* (*Zm00001eb202100*) that increase phosphorus remobilization. We also identified an upregulated set of enzymes involved in the process of substituting phospholipids with galactolipids, supported by the enrichment of glycerophospholipid metabolism pathway in KEGG (Fig 4B) and galactolipid biosynthetic process in GO (Fig 4 A), respectively. This set included the monogalactosyldia-cylglycerol synthase *mgd2* (*Zm00001eb034810*, − log_10_ *FDR* = 10.69, *log*_2_ *FC* = 4.83), the glycerophosphodiester phosphodi-esterase *gpx1* (*Zm00001eb241920*, − log_10_ *FDR* = 9.2, *log*_2_ *FC* = 6.48) and the glutathione peroxidase *glpx2* (*Zm00001eb016270*, − log_10_ *FDR* = 4.5, *log*_2_ *FC* = 7.01). Conversely, genes associated with phosphorus-intensive processes and photosynthesis were downregulated.

**Figure 4.**
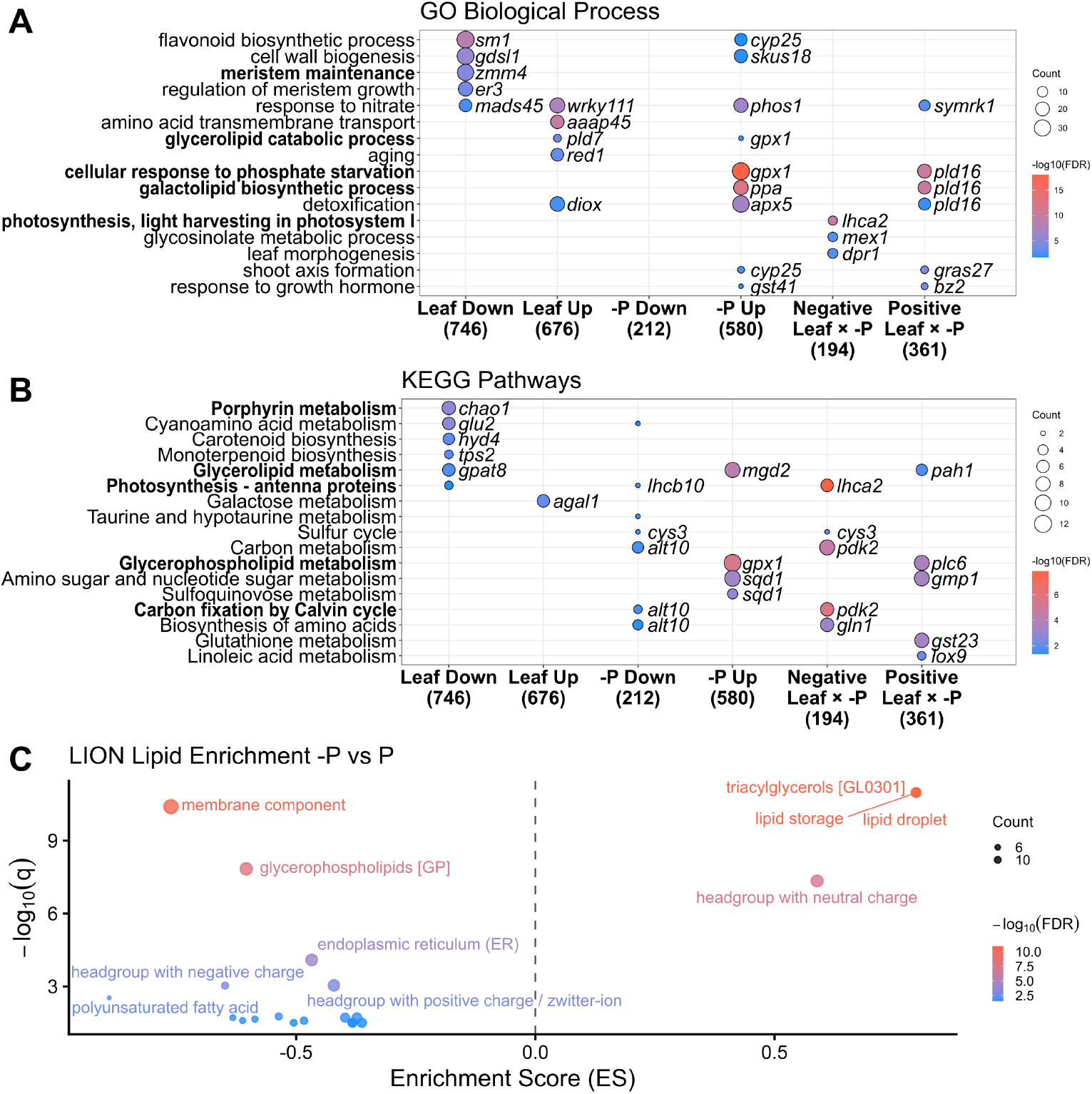
Controlled vocabulary enrichment. Overrepresentation analysis for (A) Gene ontology Biological Process, in parentheses, the total number of strong DEGs in each category, the sum of annotated genes in all overrepresented terms per category is illustrated in S6 Fig C-D, (B) KEGG Pathways, C) Lion lipids.

This includes *peamt2* (*Zm00001eb294690*, − log_10_ *FDR* = 4.93, *log*_2_ *FC* = −6.81), a phosphoethanolamine N-methyltransferase involved in phospholipid biosynthesis. Furthermore, downregulated genes consistent with reduced photosynthetic activity included light harvesting components such as *lhcb10*, and the stress responsive RUBISCO activase *rca3* (− log_10_ *FDR* = 3.8, log_2_ *FC* = −3.40). This systemic reduction in photosynthesis is also supported by the over-representation of the photosynthetic antenna proteins in KEGG, exemplified by the downregulation of the *light harvesting chlorophyll a/b binding protein10* gene (*lhcb10*) (Fig 4B). Sulfur metabolism was also affected: the chromosome 9 cysteine synthase gene cluster (*cys3*/*cys4*/*cys8, Zm00001eb376930*) was strongly repressed (− log_10_ FDR = 3.23, log_2_ FC = −1.85), driving the over-representation of the KEGG Sulfur cycle pathway (zma01320, adj. *P* = 0.023) among −P downregulated genes (Fig 4B).

Multiple transcription factors such as *zim25* (− log_10_ *FDR* = 4.2, *log*_2_ *FC* = −3.01), *nactf132* (*Zm00001eb324550*, − log_10_ *FDR* = 4.47, *log*_2_ *FC* = −4.66), and *bzip81* (− log_10_ *FDR* = 2.8, *log*_2_ *FC* = −3.37) were also repressed, suggesting a broad transcriptional reprogramming that redirects the plant resources.

### Elevated triglycerides and reduced phosphoglycerolipids are driven by phosphorus starvation and leaf age

Aggregate lipid class composition reveals the expected dominance of PC, MGDG, DGDG, and SQDG in membrane lipids, with phosphorus deficiency causing notable decreases in phos-phoglycerolipids across all leaf stages and contrasting developmental trajectories for neutral lipids under +P versus −P conditions (S8 Fig). Lipid profiling shows typical changes associated with both leaf aging and phosphorus starvation (Fig 3C-D, S7 Table, S8 Table). A multidimensional scaling of lipid abun-dance shows a marked difference between Leaf 1 and older leaves across MDS dimension 3 (17% of variance explained, Fig 3C). Older leaves were depleted in the digalactosyl diacyl-glicerol lipid DGDG36:2 ((log_2_ FC = 0.67 ± 0.15, FDR = 0.044), and accumulated the diacylglyceryl glucuronide DGGA36:3 (log_2_ FC = 0.67 ± 0.15, FDR = 0.044). The leaf age also affected the phospholipids associated with phosphatidylethanolamine (PE) turnover, as evidenced by an increase in LPC18:3 and PC36:6, and a decrease in PE32:1 (Fig 3E). Phosphorus starva-tion induces a well-characterized membrane lipid remodeling response, shifting from phosphoglycerolipids to sugar-based glycolipids. This process is illustrated by PC34:2, an abundant phospholipid that shows significant reduction in the −P main effect (log_2_ FC = −1.60 ± 0.08, FDR = 1.58 × 10^−5^) and that is further decreased in the Leaf × −P interaction (log_2_ FC = −0.56 ± 0.04, FDR = 0.0014), resulting in a decrease in concentration throughout the developmental gradient that is exacerbated by the phosphorus starvation (Fig 3E).

This widespread reduction in other phosphorus-rich mem-brane lipids is also seen in: PEs such as PE34:4, (log_2_ FC = −2.06 ± 0.27, FDR = 0.0067); Lysophosphatidylethanolamines (LPEs), with LPE18:2 being highly reduced (log_2_ FC = −2.69 ± 0.16, FDR = 7.39 × 10^−5^); and Lysophosphatidylcholines (LPCs), seen in LPC16:1 (log_2_ FC = −3.50 ± 0.30, FDR = 0.0007). Concomitantly, phosphorus starvation leads to a storage response through the accumulation of triacylglycerols (TAGs), evidenced by the highly upregulated TG56:6 (log_2_ FC = 12.71 ± 0.62, FDR = 0.018) (Fig 3D) LION lipid enrichment analysis confirms these systemic changes, showing an extremely strong enrichment of triacylglycerols (FDR = 1.06 × 10^−11^, ES = 0.80) and associated lipid storage terms, alongside a highly significant decrease in glycerophospholipids (FDR = 1.46 × 10^−8^, ES = −0.60) and membrane components (FDR = 4.03 × 10^−11^, ES = −0.76) (Fig 3C) We did not detect strong DALs for the *Inv4m* main effect.

### Phosphorus starvation promotes senescence-associated transcription and the shutdown of light-dependent photosynthesis reaction genes

To understand the interplay between leaf development and nutrient stress, we first established the baseline transcriptional signatures of leaf aging. We observed significant, opposing correlations between leaf stage and the expression of key biological processes (Fig 5. Chlorophyll Synthesis shows a significant negative correlation with age (*r* = −0.78, *p* = 6.3 × 10^−10^), while Chlorophyll Degradation shows a positive correlation (*r* = 0.53, *p* = 2.5 × 10^−4^). Photosynthesis shows a significant negative correlation (*r* = −0.76, *p* = 2.6 × 10^−9^), and Leaf Senescence shows a significant positive correlation (*r* = 0.77, *p* = 1.2 × 10^−9^)). For the chlorophyll index, downregulation with leaf age was driven by consistent decreases in *chlh1* (*Zm00001eb433610*, Mg chelatase subunit H 1), *gtr3* (*Zm00001eb044210*, glutamyl-tRNA reductase), and *urod* (*Zm00001eb358510*, uroporphyrinogen decarboxylase), while the degradation enzyme *pph* (*Zm00001eb231810*, pheophytinase) showed the opposite pattern (Fig 5 B). For the photosynthesis and senescence indices, the trajectories were exemplified by declining expression of *pep1* (*Zm00001eb383680*, phosphoenolpyruvate carboxylase) and *ssu1* (*Zm00001eb197410*, RuBisCO small subunit) concurrent with upregulation of *salt1* (*Zm00001eb407630*) and *mir3* (*Zm00001eb068400*) (Fig 5 B). Notably, the STAY-GREEN homologs *sgrl1* (*Zm00001eb076680*) and *nye2* (*Zm00001eb103480*) exhibited opposing expression patterns despite similar functional annotations, suggesting divergent roles in senescence regulation (Fig 5 B) Phosphorus deficiency significantly amplified these developmental programs, generating 487 genes with significant leaf stage × phosphorus interactions (Fig 5 C, E). Under −P conditions, the divergence between anabolic and catabolic processes intensified with leaf age: chlorophyll degradation (*p <* 0.01), chlorophyll synthesis (*p <* 0.05), photosynthesis (*p <* 0.001), and senescence indices (*p <* 0.01) all showed significant interactions between developmental stage and nutrient status (Fig 5D). Genes with negative interaction terms, including and *tat, Tat pathway signal sequence family protein* (*Zm00001eb359280*) and *cab, chlorophyll a-b binding protein* (*Zm00001eb207130*), showed larger negative P-starvation responses in older leaves, while those with positive interactions such as *pk, pyruvate kinase* (*Zm00001eb157810*) and *mrpa3, multidrug resistance protein* (*Zm00001eb376160*), which is involved in anthocyanin transport into the vacuole (Goodman *et al*. 2004), displayed amplified positive responses in older leaves (Fig 5 F). KEGG pathway enrichment in the Leaf:−P downregulated cluster was strongest for Photosynthesis antenna proteins (zma00196, adj. *P* = 1.4 × 10^−8^; exemplified by *lhca2* and *lhcb10*) and Carbon fixation by Calvin cycle (zma00710, adj. *P* = 1.6 × 10^−6^; exemplified by *pdk2*), indicating that both light harvesting and carbon fixation capacity are progressively shut down in older leaves under −P (Fig 4B). The Sulfur cycle pathway (zma01320, adj. *P* = 9.3 × 10^−3^) remained enriched in the Leaf:−P downregulated cluster through the continued repression of the *cys3*/*cys4*/*cys8* gene cluster with leaf age, while the Leaf:−P upregulated cluster was enriched for Glutathione metabolism (zma00480, adj. *P* = 3.5 × 10^−4^), driven by a panel of glutathione S-transferases including *gst23*. This interaction pattern indicates that phosphorus deficiency not only triggers immediate metabolic adjustments but also accelerates the natural developmental program of leaf senescence, with older leaves experiencing disproportionately severe molecular stress responses that compound the effects of nutrient limitation.

**Figure 5.**
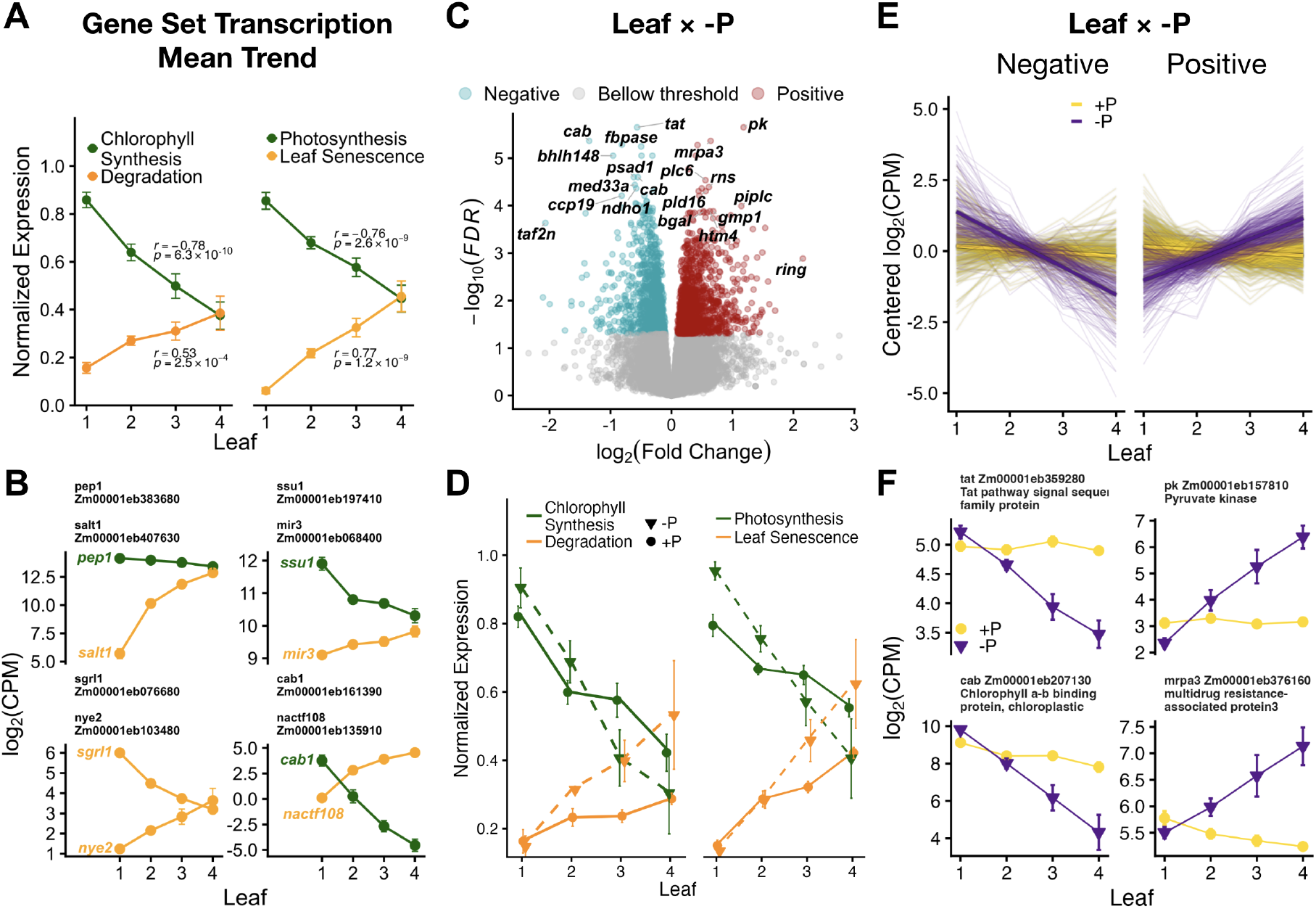
The response to phosphorus starvation increases with leaf stage and is positively correlated with indicators of leaf senescence. (**A**) Gene Set Transcription Indices Across Leaf Stages. Indices are calculated as the mean log_10_(CPM) for genes within defined sets and normalized across the four leaf stages to represent the proportion of the total expression range. The left panel shows Chlorophyll Synthesis (dark green) and Chlorophyll Degradation (light green/orange) sets derived from CornCyc/KEGG; the right panel shows Photosynthesis (dark green) and Leaf Senescence (orange) sets from (Ojeda-Rivera *et al*. 2026). (**B**) Expression profiles (log_10_(CPM)) for representative gene pairs illustrating opposing trends across leaf stages. Error bars represent SEM. (**C**) Volcano Plot of Leaf × Phosphorus (P) Interaction. The plot highlights genes with a significant transcriptional interaction between leaf stage and phosphorus treatment (+P vs. −P). Genes with a negative log_2_(Fold Change) and significant FDR are colored red (negative interaction), and those with a positive log_2_(Fold Change) and significant FDR are colored blue (positive interaction). (**D**) Gene Set Transcription Indices Split by Phosphorus Treatment. The mean normalized expression for Chlorophyll Synthesis/Degradation (left) and Photosynthesis/Senescence (right) is plotted for phosphorus-deficient (−P, dashed lines) and phosphorus-sufficient (+P, solid lines) conditions. (**E**) Individual Gene Trajectories for Leaf × Phosphorus Interaction. Genes are partitioned based on their interaction term (Negative: left, Positive: right). Thin lines represent individual gene expression profiles (centered log_2_(CPM)) across the four leaf stages under −P (purple) and +P (yellow) conditions. Bold lines illustrate the mean trend for each group. (**F**) Expression profiles (log_10_(CPM)) for representative genes from the interaction set. *Right*: leaf × P interaction (e.g., *multidrug resistance protein mrpa3* and a *Chlorophyll a-b binding protein*). Error bars represent SEM.

## Discussion

From our multi-omics analysis, we can infer that the maize phosphorus starvation response is shaped by leaf developmental stage, with older leaves showing enhanced stress responses indicative of the onset of developmental senescence during the vegetative phase. While phosphorus deficiency triggered canonical molecular responses across genotypes, the magnitude of these responses varied depending on the leaf developmental position.

### We captured a gradient of sequential leaf senescence in our samples

By sampling leaves from the topmost fully developed collar and those below, we captured a physiological gradient in gene expression. The gradient appears to accurately reflect the onset and initial phases of sequential senescence in our plants, which we estimated around the V13 stage, approximately 10 to 16 days prior to flowering. We observed this sequential senescence as the progressive aging of the leaves along the plant axis, culminating in the death of the leaves nearest to the soil.

Using gene set transcriptomic indices, we corroborated that chlorophyll biosynthesis/degradation and photosynthesis/senescence were correlated with this vertical developmental axis Fig 5 A. In the chlorophyll catabolism pathway, *pph* showed positive correlation with leaf age while *sgrl1* was downregulated, potentially shifting chlorophyll catabolism towards PPH from the chlorophyllase pathway, which has been reported to mediate 87% of chlorophyll degradation in maize (Wei *et al*. 2025).

This developmental framework provides context for interpreting phosphorus starvation responses. The leaf-stage variable represents a combined axis of decreasing photosynthetic capacity and the progression of developmental senescence, allowing us to quantify how nutrient stress interacts with natural developmental transitions during vegetative growth. The significant interaction terms we observed for chlorophyll metabolism, photosynthesis, and senescence gene sets indicate that phosphorus deficiency does not uniformly shift all leaves along this gradient but rather amplifies the divergence between young and old tissues Fig 5D.

### Phosphorus starvation responses accelerate with leaf age

Phosphorus deficiency activated the expected molecular machinery for nutrient scavenging and remobilization. Upregulated genes included non-coding RNAs *pilncr1* and *ips1*, phosphate scavenging enzymes *gpx1, pap2* (*Zm00001eb064450*), and *pap19*, phosphate transporters *phos1, pht1*, and *pht7*, and galactolipid biosynthesis genes *mgd2, sqd2* (*Zm00001eb297970*), *sqd3* (*Zm00001eb335670*), and *glpx2*. Gene Ontology enrichment confirmed activation of galactolipid biosynthetic processes and phosphate starvation response pathways, while KEGG analysis highlighted glycerophospholipid metabolism (Fig 4 A-B). Concurrent with this phospholipid to galactolipid swap, transcriptome data show induction of *pah1* (phosphatidate phosphatase, log_2_ FC = +1.30) and *dgk6* (diacylglycerol kinase, log_2_ FC = +0.46), which produce the diacylglycerol substrate for galactolipid synthesis, alongside repression of phospholipase D paralogs (*pld10, pld13, pld15*) and a chloroplastic Phospholipase A1. Downregulated genes included *peamt2* (*Zm00001eb294690*), which catalyzes phosphoethanolamine methylation in the Kennedy pathway for PC biosynthesis, the light harvesting gene *lhcb10* (*Zm00001eb357740*) and the stress responsive RUBISCO activase *rca3* (*Zm00001eb164380*), and transcription factors *zim25* (*Zm00001eb278320*), *nactf132* (*Zm00001eb324550*), and *bzip81* (*Zm00001eb198410*). These patterns validate the canonical phosphorus starvation response documented in previous studies (Wang *et al*. 2020; He *et al*. 2022).

We detected 555 strong DEGs with a significant and strong interaction between leaf stage and phosphorus treatment Fig 4A-B, meaning that their phosphorus responses increased or decreased linearly with leaf position. Our statistical modelling was designed to detect two functionally distinct trajectories. Genes with negative interaction terms (194), including light harvesting proteins *cab, psad1*, and *ndho1*, showed increased negative responses to phosphorus starvation in older leaves, corresponding to selective shutdown of the thylakoid light-capture machinery. Gene Ontology enrichment for this set highlighted terms related to photosynthesis and the light harvesting complex, and KEGG analysis identified both Photosynthesis – antenna proteins (zma00196) and Carbon fixation by Calvin cycle (zma00710) as strongly enriched in the Leaf:−P downregulated cluster, with both pathways gaining substantially more DEGs in the interaction cluster than in the −P main effect cluster (Fig 4A-B). Both light harvesting and carbon-fixation capacity are therefore progressively shut down in older leaves under phosphorus limitation, pointing to a coordinated decommissioning of the photosynthetic machinery rather than a split between components that are uniformly regulated across leaf stages and components whose regulation is amplified with leaf age.

Genes with positive interaction terms (Fig 4A-B) i.e. with increased slope of *log*_2_FC *>* 0.5 per leaf stage under phosphorus limitation, showed amplified phosphorus-starvation responses in older leaves, corresponding to enhanced senescence and nutrient remobilization signatures. Out of 361 strong DEGs with positive interaction with leaf age, 20 genes were annotated with “cellular response to phosphate starvation” and presented in the S6 Table. These genes can be classified into more specific functional groups. The glycolytic enzymes, *pfk1*, together with *pep2* and its activating kinase *ppck3*. The galactolypid byiosynthetic enzymes *mgd3, sqd2*, and *sqd3*, were strongly upregulated, indicating accelerated replacement of phospholipids with sulfo-and galactolipids in ageing plastids. The SPX-domain proteins *spx2* and *spx6*, central regulators of the phosphate-starvation signalling cascade, showed the same trajectory, confirming that the sensing machinery itself is amplified in older tissue. Phosphorus-scavenging capacity was reinforced by pap1 and the three inorganic pyrophosphatase paralogues (*ppa1, ppa2, ppa3*) that recycle Pi from cytosolic pyrophosphate. Phospholipid turnover was evidenced by leaf age-dependent upregulation of *pld16* and the ER-associated *pah1*. Collectively, these age-amplified responses reveal a programmed shift from phosphorus conservation to wholesale phosphorus remobilization as maize leaves senesce under phosphorus limitation.

### PC34:2 depletion increases with leaf age and correlates with flowering delay and the downregulation of flowering transcription factors

Lipidomic analysis confirmed the classical membrane remodeling response to phosphorus starvation. Phospholipid classes showed widespread depletion: phosphatidylcholines PC34:2, PC32:2, PC32:0, and PC38:6; phosphatidylethanolamines PE34:4, PE34:3, and PE32:1; lysophosphatidylethanolamines LPE18:2, LPE18:3, and LPE16:0; lysophosphatidylcholines LPC16:1, LPC18:3, and LPC18:2; and phosphatidylglycerols PG32:0 and PG34:3. Triacylglycerols accumulated under phosphorus stress, with TAG species TG50:3, TG52:6, TG54:9, TG56:6, TG50:2, and

TG52:3 showing increased abundance (Fig 3E). TG56:6 exhibited accumulation exceeding 12-fold increase. Galactolipid accumulation was evidenced by DGGA36:4, a phosphorus-free membrane component (Fig 3E). LION lipid enrichment analysis confirmed these systemic changes: triacylglycerols showed strong enrichment, while glycerophospholipids and membrane components were significantly depleted (Fig 4C).

These patterns align with the established membrane remodeling strategy documented previously: replacement of phosphorus-containing phospholipids with galactolipids conserves phosphorus for nucleic acid and ATP synthesis, while TAG accumulation provides temporary storage for fatty acids released from membrane degradation (Wang *et al*. 2020).

Phosphatidylcholine PC34:2 showed both a strong main effect of phosphorus deficiency and a significant Leaf × −P interaction, indicating that the magnitude of PC34:2 depletion increased systematically with leaf developmental position (Fig 3E). This age-dependent response is noteworthy given PC34:2’s binding to ZCN8, the maize florigen ortholog (Barnes *et al*. 2022). We previously demonstrated that PC34:2 copurifies with recombinant ZCN8 protein and identified probable binding sites through molecular docking simulations (Barnes *et al*. 2022). Additionally, phospholipase HPC1 expression, which influences PC34:2 levels, correlates with flowering-time variation across the maize diversity panel (Barnes *et al*. 2022).

While *Inv4m* does not modulate the PC34:2 response, the coordinate patterns we observed (PC34:2 depletion, flowering delay under phosphorus stress, and *peamt2* suppression) raise the possibility that phosphorus stress influences reproductive timing through multiple mechanisms. The dramatic suppression of *peamt2*, which catalyzes sequential methylation of phosphoethanolamine to phosphocholine in the Kennedy pathway, represents one of the strongest transcriptional responses in our dataset. In Arabidopsis, the ortholog XIPOTL1 affects flowering time, root architecture, and stress responses (Cruz-Ramírez *et al*. 2004). In maize, natural variation at the *peamt2* locus associates with flowering time in Mexican highland populations (Perez-Limón *et al*. 2022; Barnes *et al*. 2022), suggesting that modulation of phospholipid biosynthesis contributes to adaptive variation in developmental timing. If ZCN8 florigen activity or stability requires PC34:2 binding, as demonstrated for *Arabidopsis* FT interactions with phosphatidylglycerol in temperature-dependent contexts (Susila *et al*. 2021), then systematic depletion of this specific phospholipid species could potentially impair florigen signaling beyond resource limitation alone.

The downregulation of MADS-box transcription factors *zmm4* and *zmm15* with increasing leaf age further connects lipid remodeling to developmental transitions, as these two transcription factors are associated with the highest flowering time in a TWAS analysis of the Wisconsin panel (Torres-Rodríguez *et al*. 2024). These flowering-time regulators showed strong negative correlations with leaf stage, consistent with the natural progression from vegetative to reproductive phase transitions during our sampling window around V13 stage.

### LPE is a senescence marker in our maize samples despite its anti-senescence role in tomato and Arabidopsis

16:3 plants (*Arabidopsis*, tomato, spinach) and 18:3 plants (maize and most grasses) differ in how their chloroplast monogalactosyldiacylglycerol (MGDG) is assembled: in 16:3 plants a substantial fraction of MGDG is built *in situ* through the plastid (“prokaryotic”) pathway, which preserves a 16-carbon fatty acid at the sn-2 position; in 18:3 plants MGDG is built predominantly from ER-derived (“eukaryotic”) precursors imported back into the plastid, placing an 18-carbon fatty acid at sn-2. Both pathways operate in all plants; the terminology reflects the dominant route. It has been demonstrated in 16:3 plants such as *Arabidopsis* and tomato, that LPE acts as a leaf senescence suppressor (Amaro and Almeida 2013). In these plants, the application of exogenous LPE delays senescence by inhibiting phospholipase D and preserving membrane integrity (Amaro and Almeida 2013; Ryu *et al*. 1997).

The behavior of lysophosphatidylethanolamines in our dataset contradicts this established pattern, highlighting a possible new difference in phospholipid metabolism between 16:3 and 18:3 plants. Although multiple LPE species (LPE18:2, LPE18:3, and LPE16:0) were depleted under phosphorus deficiency, LPE18:3 showed a strong positive association with increased leaf developmental stage. This accumulation in our samples suggests that, at endogenous concentrations, LPE functions as a marker for initial senescence, contrary to its reported senescence-suppressor role in 16:3 plants.

The lipidomic pattern is paralleled by coordinated transcriptional regulation. Phospholipases *plc6* and *pld16* both showed positive Leaf × −P interactions, indicating enhanced phospholipid hydrolysis in older leaves under phosphorus stress. In contrast, lysophosphatidylethanolamine acyltransferase *lpeat2* and choline/ethanolamine kinase *cek4* showed no differential expression, suggesting constitutive regulation.

This divergence might reflect the difference between 16:3 and 18:3 plants in lipid metabolism architecture (Mongrand *et al*. 1998; Heinz and Roughan 1983). Maize GDG has 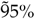 C18/C18 species thought to derive from the eukaryotic pathway of GDG biosynthesis. Previous reports indicated that maize was devoid of C18/C16 GDG species (Mongrand *et al*. 1998), however, the use of mass spectrometry now reveals that a minor amount of GDG biosynthesis proceeds through the prokaryotic pathway in both maize leaves and endosperm (Myers *et al*. 2011). In maize’s eukaryotic pathway, diacylglycerol derived from phospholipid degradation is transported to the chloroplast outer envelope for galactolipid synthesis, creating direct metabolic flux from PE to LPE to DAG to MGDG (Gu *et al*. 2017). The efficiency of this pathway, particularly when PC biosynthesis is blocked by *peamt2* suppression, may preclude LPE accumulation as the lipid is rapidly consumed in downstream reactions. This positions LPE as a transient salvage intermediate rather than a regulatory signal in 18:3 plants.

### Enhanced PAH1 activity, but not PDAT expression, underlies leaf stage-dependent accumulation of polyunsaturated TAGs

The accumulation of triacylglycerols (TAGs) in phosphorus-deficient leaves might reflect a substrate-driven lipid salvage mechanism during membrane remodeling. Phosphorus starvation led to enrichment of highly unsaturated TAG species, including TG(54:9) and TG(52:6), both containing 18:3 fatty acids, coinciding with depletion of polyunsaturated phospholipids such as PC(36:6) and LPC(18:3). The preservation of polyunsaturated fatty acids in TAG form is consistent with direct transfer via the PDAT pathway, which moves intact acyl chains from phospholipids to diacylglycerol (Chen and Smith 2012), avoiding the energetic cost of complete phospholipid degradation and subsequent re-desaturation. However, TAG accumulation occurred without significant transcriptional up-regulation of PDAT genes, indicating that the phospholipid-to-TAG flux is driven primarily by substrate availability rather than increased PDAT expression.

Phosphatidate phosphatase *pah1* showed a positive Leaf × −*P* interaction (+0.58 log_2_ FC per leaf stage), indicating higher expression in older leaves under phosphorus stress. This increase in activity may be directing lipid flux toward TAG synthesis versus PC synthesis in our samples, as observed in *Arabidopsis* (Eastmond *et al*. 2010). Notably, *pah1* and the co-upregulated phospholipase *pld16* (+0.56 log_2_ FC per leaf stage) are orthologous to *Arabidopsis* PAH1 and PLD*ζ*2, which act together in a pathway to convert phospholipid-derived phosphatidic acid into diacylglycerol for TAG assembly during phosphorus deficiency. Beyond lipid remodeling, PLD*ζ*2 has been linked to autophagy and vacuolar acidification (Guan *et al*. 2025), suggesting coordination between membrane catabolism and cellular recycling to supply lipid substrates under phosphorus stress.

These findings indicate that, in our samples, phosphorus-dependent TAG accumulation is primarily driven by enhanced PAH1-mediated flux from phospholipids to diacylglycerol rather than by PDAT expression. This substrate-driven mechanism supports transient storage of fatty acids released from phospholipid catabolism, either for transport to younger tissues or for energy production via *β*-oxidation (Wang *et al*. 2020). The age-dependent amplification of TAG accumulation in our dataset is compatible with this interpretation and suggests that substrate-driven lipid salvage operates in senescing tissues during the vegetative phase.

### Leaf senescence transcription accelerates under phosphorus limitation

The coordination of developmental senescence with phosphorus starvation might be an adaptive strategy in maize, where leaf age influences the magnitude of nutrient stress responses. Five *Arabidopsis* ortholog genes (SAGs) in (Zhang *et al*. 2014b) are strong DEGs for the effect of leaf stage, providing support for our observed leaf developmental gradient. The NAC transcription factor *nactf108* (*Zm00001eb135910*, ORE1/ANAC092), a regulator known to activate SAG targets during early senescence, was found to increase with leaf position. Similarly, transcripts for chlorophyll degradation enzymes *pph* (*Zm00001eb231810*) (pheophytinase) and *nye1* (*Zm00001eb319560*,SGR/STAY-GREEN), along with proteolytic machinery including *see2b* (*Zm00001eb162210*, gamma vacuolar processing enzyme) and *clpb1* (*Zm00001eb242420*, ERD1/SAG15), progressively increased from young to old leaves. This expression pattern is consistent with the canonical progression of natural leaf aging during vegetative growth, establishing a baseline for interpreting stress-induced deviations. A key observation in our study is the Leaf × −P interaction, which indicates that phosphorus stress does not uniformly affect all leaves but rather compounds with leaf age to create stronger responses in older tissues (Fig 5)

The ribonuclease *rns* (*Zm00001eb144680*) exemplifies this pattern, exhibiting both a strong main effect of phosphorus starvation and significant age-dependent amplification, which suggests that RNA degradation may accelerate specifically where developmental senescence and nutrient stress converge. This interaction extends to *csap* (*Zm00001eb402430, chloroplast-localized senescence-associated protein*), implying a potential co-ordination in the dismantling of the photosynthetic apparatus when both stressors are present, (So *et al*. 2020). We identified 24 genes within this Leaf × −P interaction term that reveal potential functional specialization. Three hub genes, *rns, mybr105* (*Zm00001eb081290*, a protein-binding MYB), and *lkrsdh1* (*Zm00001eb192910*) (involved in lysine catabolism), appear central to these nutrient salvage operations, while transport genes (e.g., the carbohydrate transporter *sweet2* (*Zm00001eb342040*) and *ppt1* (*Zm00001eb097690*) and cell wall remodeling enzymes, *aga2* (*Zm00001eb281720*) and *irx15* (*Zm00001eb068410*), are upregulated, consistent with nutrient remobilization.

To map the regulatory architecture governing these processes, we compiled a list of SAGs by cross-referencing our strong DEGs with manually curated sources (Zhang *et al*. 2014b; Liu *et al*. 2011; Ojeda-Rivera *et al*. 2026; Berardini *et al*. 2015; Durinck *et al*. 2005), resulting in a total of 110 genes (S1 File). Almost half of these, 53, were transcription factors. They are distributed across 13 families, with the NAC (16 members), WRKY (7), G2-like (5), AP2-EREBP (5), MYB (5), and bHLH (5) families representing the dominant regulatory nodes. Within the NAC family, we observed complex, context-dependent expression patterns. Typical senescence regulators like *nactf108* and *nactf44* (*Zm00001eb015630*) consistently increase with leaf age, and *nactf132* (*Zm00001eb324550, ZmNAC132*) shares this leaf-age upregulation while also showing strong downregulation under phosphorus stress. This divergence is particularly noteworthy because *nactf132* has been linked to chlorophyll content regulation through *nye1* activation (Yuan *et al*. 2023), and natural variation in its 5^′^UTR associates with chlorophyll B levels (Wallace *et al*. 2014). The contrasting responses suggest that developmental senescence and nutrient-stress-amplified senescence might engage partially distinct regulatory networks despite converging on common downstream targets.

Beyond the NAC family, other regulatory groups show similar complexity. The Leaf × −P interaction specifically captures *nactf32* (*Zm00001eb080700*), which shifts from downregulation in standard aging to upregulation under combined stress. The WRKY family also contributes to this stress-integrated network, with *wrky17* (*Zm00001eb330710*), *wrky32* (*Zm00001eb015320*), and *wrky92* (*Zm00001eb350280*) specifically responding to the age-phosphorus interaction. The AP2-EREBP and MYB families also contribute to this stress-integrated network, with members like *myb112* (*Zm00001eb387370*) and *myb163* (*Zm00001eb366540*) responding strongly to phosphorus limitation, potentially bridging the gap between metabolic signaling and transcriptional control. Our analysis suggests that phosphorus limitation may accelerate natural senescence programs specifically in older leaves through transcriptional cascades that integrate developmental timing with nutrient availability.

The observed top-bottom gradient of responses suggests that phosphorus deficiency accelerates rather than replaces natural senescence programs. Senescence-associated gene expression increases with age, phospholipids deplete, triacylglycerols accumulate, and photosynthesis declines. All of these responses under phosphorus stress are amplified and propagated basipetally (downwards) along the plant axis during the vegetative phase. At the extreme bottom, this culminates in the senescence and sacrifice of lower-canopy leaves to remobilize nutrients, thereby sustaining younger, photosynthetically active tissues and developing reproductive structures (Wei *et al*. 2025). Even the observed increase in the seed-to-stover phosphorus ratio under deficiency supports this concept of prioritized allocation.

We believe that crop improvement strategies might benefit from focusing on optimizing the timing and tissue specificity of senescence responses.

### Phosphorus starvation reshapes mineral partitioning between stover and grain

Under standard fertilization, maize already partitions nearly 80% of total phosphorus to grain through a combination of uptake after silking and remobilization of leaf and stalk reserves bender2013,veneklaas2012. In our data, −P reduces stover phosphorus concentration far more than seed phosphorus concentration, nearly doubling the seed/stover ratio (Fig 2). Because we measured tissue concentrations rather than content per plant, we cannot separate how much of this shift reflects reduced stover uptake versus increased remobilization from stover to grain. These two fluxes contribute to grain phosphorus in varying proportions across genotypes under low P sun2023,roller2025.

The seed/stover ratios shift in opposite directions for phosphorus and zinc under −P, against a background of higher sulfur in both tissues (Fig 2). These shifts are consistent with the established shoot ionomic signature of phosphorus homeostasis baxter2008 and the phosphate-sulfate and phosphate-zinc cross-talk described in *Arabidopsis* and maize bielecka2014,bouain2014,wangzou2023,roller2025. For zinc in particular, the canonical high harvest index (~ 62%) and ~ 60% re-mobilization from stalk to grain observed under standard fertilization bender2013 appear inverted under −P, with zinc retained in source tissues at the expense of grain loading. Direct measurements of increased shoot zinc under phosphorus limitation have been documented in maize soltanghei2014,saenchai2016, sorghum oseni2009, finger millet maharajan2023, and *Arabidopsis* under long-term Pi deprivation misson2005. In contrast, the molecular dissection of Pi-Zn crosstalk has been performed primarily in the reverse direction, with Zn deficiency driving shoot Pi over-accumulation through the master phosphate starvation transcription factor PHR1 bouain2014,khan2014; whether PHR1 also mediates the zinc retention observed here under phosphorus limitation remains to be tested. In our data, the PHR1 paralog *glk36* (*PHL11, Zm00001eb071920*, log_2_ FC = +1.53) and the SPX-domain sensor *spx2* (log_2_ FC = +4.90) are strongly induced under −P, consistent with active PHR1 signalling. Calcium and magnesium showed smaller shifts; grain calcium concentrations in modern maize are too low to be quantified reliably in nutrient partitioning analyses bender2013, and we therefore interpret the calcium and magnesium directions as descriptive.

The transcriptional data recover the sulfolipid arm of this crosstalk, with *sqd1, sqd2* and *sqd3* among the strongest −P upregulated genes, consistent with SQDG biosynthesis upregulation seen across maize genotypes he2022. The chromosome 9 cysteine synthase gene cluster (*cys3*/*cys4*/*cys8* at *Zm00001eb376930*, and the adjacent *cys7*) is repressed under −P with the repression deepening in older leaves (Leaf×−P interaction), and the Sulfur cycle pathway (KEGG zma01320) is overrepresented in both the −P and Leaf:−P downregulated clusters at this locus. The cys3/cys4 product physically interacts with the E3 ubiquitin ligase NLA in yeast two-hybrid assays liao2026, a regulator of phosphate homeostasis that drives ubiquitination and turnover of PHT1 transporters together with the E2 conjugating enzyme PHO2 huang2013, raising the possibility of a post-translational link between cysteine biosynthesis and Pi transporter regulation beyond the transcriptional repression observed here. Downstream glutathione machinery moves in the opposite direction: *γ*-glutamylcysteine synthetase *gsh1* is a Leaf×−P up-regulated gene, and Glutathione metabolism (KEGG zma00480,adj. *p* = 3.5 × 10^−4^) together with the glutathione metabolic process (GO:0006749) are enriched among Leaf:−P upregulated genes, driven by a panel of glutathione S-transferases. In older −P leaves, the transcriptional program for sulfolipid biosynthesis (*sqd1, sqd2, sqd3*) is induced and the glutathione utilization machinery is progressively amplified, while new cysteine synthesis at the dominant chromosome 9 locus is reduced.

### Inv4m does not alter phosphorus starvation responses

Across whole plant phenotypes, the leaf transcriptome, lipidome, and ionome, phosphorus deficiency resulted in consistent starvation responses. However, we did not find decisive evidence that the *Inv4m* inversion modulates any of them. The two exceptions were incidental: a significant *Inv4m* × phosphorus interaction on cob diameter (Fig. 1F) and on the expression of *aldh2* (S7 Fig), a gene located in the flanking introgression of the inversion and without annotation related to phosphorus. No candidate gene for the phosphorus starvation response residing within the inversion showed a significant *Inv4m* ×−P interaction at the transcriptional level. This includes *phos2* (*Zm00001eb191650*, ZmPHO1;2a), a PHO1 family phosphate efflux transporter and a candidate for adaptation to low phosphorus availability within the inversion (Salazar-Vidal *et al*. 2016; **?**; **?**; **?**).

Nonetheless, we detected additive *Inv4m* effects on the ionome, most prominently reductions in stover Ca, Zn, and K content and differences in Fe grain content and harvest index (S5 Fig). However, *Inv4m* plants showed lower stover biomass at harvest (marginal Inv4m vs CTRL contrast: −7.9 ± 3.3 g STWHV, FDR = 0.044), so the stover content differences may in part reflect this biomass difference rather than a distinct mineral reallocation mechanism.

The *Inv4m* phenotypic effects on flowering time, plant height, and related traits are developed in detail in the companion manuscript (Rodriguez-Zapata *et al*. 2026).

## Conclusion

Our multiomics analysis reveals that the maize phosphorus starvation response is influenced by the leaf developmental stage during the vegetative phase, with older leaves positioned below the collar exhibiting enhanced stress responses characteristic of developmental senescence, which integrate nutrient limitation with natural developmental progression. The bifurcation of molecular responses into light harvesting shutdown and senescence acceleration shows coordinated regulation of functionally distinct pathways. Lipidomic patterns parallel transcriptomic responses, with age-dependent amplification of phospholipid degradation, galactolipid accumulation, and triacylglycerol synthesis. The divergence of lysophosphatidylethanolamine patterns from 16:3 plant models highlights the importance of lipid metabolism architecture differences between 16:3 and 18:3 plant species. Despite this strong developmental dependency, the *Inv4m* chromosomal inversion does not substantially modulate phosphorus starvation responses, indicating that its contribution to highland adaptation operates through constitutive effects on developmental timing rather than enhanced nutrient stress tolerance.

## Materials and methods

### Inv4m Near Introgressed Lines, growth conditions, experimental design, and phenotype measurements

To measure the effects of *Inv4m* in plant field phenotypes and their phosphorus starvation response transcriptome, we used a highland traditional variety carrying the Highland haplotype of *Inv4m* corresponding to the inverted karyotype. The accession Michoacán 21 (referred to as Mi21), from the Mexican Cónico group, was obtained from the International Maize and Wheat Improvement Center (CIMMYT). In contrast, the reference genome of the temperate inbred B73, the recurrent parent for introgression, carries the lowland haplotype corresponding to the standard non-inverted karyotype at *Inv4m*. From the cross of Mi21 with B73, one F1 individual was backcrossed to B73 for eight generations. We selected lines carrying *Inv4m* with a diagnostic SNP during each cycle using a cleaved amplified polymorphic sequence (CAPS) marker. The marker SNP is PZE04175660223 located at position 4:181637780 in the NAM B73v5 *Zea mays* genome assembly. Amplification of the polymorphic site was done with the following primer pair: *CTGAGCAGGAGATGATG-GCCACTC* and *GGAAAGGACATAAAAGAAAGGTGCA*, and subsequently cleaved by *HinfI*. Plants were genotyped using the CASP marker for selecting heterozygous plants at BC8S2 after selfing seeds of *Inv4m* and CTRL homozygous individuals were selected for the field trial.

Plants were planted on May 26 2022 at the Russell E. Larson Agricultural Research Farm in Rock Springs, Pennsylvania (40°42’36” N 77°57’0” W, 366 m.a.s.l.) in soil classified as a Hagerstown silt loam (fine, mixed, semiactive, mesic Typic Hapludalf). Experimental conditions were similar to previously described (Strock *et al*. 2018). The experiment had a complete block design with two phosphorus (P) levels. Low-P fields (5 ppm Melich-3 Phosphorus) and high-P fields (36 ppm Melich-3 Phosphorus) were divided into smaller blocks. Three rows per block were planted, with a mean stand count of 8 plants per plot. The plants from the center row were selected for measurements to minimize border effects. Fields received fertilization based on treatment requirements. Drip irrigation was provided during dry periods. Each genotype was replicated four times within its P treatment.

### Inductively Coupled Plasma Optical Emission Spectroscopy (ICP-OES)

Stover samples were dried and homogenised at Pennsylvania State University; both stover and seed samples were shipped to the USDA-ARS Robert W. Holley Center for Agriculture and Health, Cornell University, for elemental analysis by Inductively Coupled Plasma Optical Emission Spectroscopy (ICP-OES). Samples were weighed, digested in double-distilled HNO_3_ followed by a 60/40 nitric/perchloric acid mixture at 150 °C, and diluted to a final volume of 10 mL with deionised water prior to analysis on a Thermo iCap 7000 ICP-OES (**?**). Element concentrations were reported per sample dry weight. Signals for Al, B, Ca, Fe, K, Mg, Mn, Mo, Ni, P, S, Zn, Na and Cu were quantified; values beyond three times the interquartile range were flagged as outliers and excluded from downstream analysis.

Ionome responses were modeled with the same two-stage linear-model framework described for the phenotypes below (Eq. 1), fit independently to 48 mineral × metric combinations covering 8 minerals (P, Zn, Ca, S, Mg, Mn, K, Fe) and 6 metrics per mineral (concentration in stover and grain, grain/stover concentration ratio, content in stover and grain, mineral harvest index). Within-genotype −P vs +P ionome contrasts formed a single emmeans (**?**) family of 96 tests (48 responses × 2 within-genotype contrasts) and are plotted for P, Zn, Ca, S in Figure 2 and for Mg, Mn, K, Fe in Supplementary Figure S4 Fig. Marginal *Inv4m* vs CTRL ionome contrasts pooled across phosphorus treatments formed a separate family of 48 tests; Supplementary Figure S5 Fig plots this contrast for Mg, Ca, Zn, Fe, and K. Both families were adjusted with Benjamini-Hochberg FDR and reported at *FDR <* 0.05.

### Phenotype analysis

For stover mass growth curves, a different plant at each time point (40, 50, 60, and harvest at 121 days after planting) was collected, dried, and weighed for each row. Stover dry mass data was fitted to a logistic growth model using the R package *Growthcurver* sprouffske2016. Maximum stover dry weight was estimated as the maximum across the four time points and not dry weight at harvest. Ear measurements were taken for one ear per row at harvest.

We used a two-stage approach to model phenotypic responses to phosphorus treatment and genotype. In the first stage, we applied spatial correction separately for each treatment (+P and −P) using P-spline analysis of spatial trends (SpATS) rodriguez-alvarez2018 implemented via the `statgenHTP::fitModels()` function millet2025. For each phenotype *y* and treatment separately, SpATS fits a mixed model that decomposes spatial variation into smooth bivariate surfaces using two-dimensional P-splines over field row and column co-ordinates, while accounting for genotypic effects and replication structure. The model estimates genotype-specific predictions adjusted for spatial heterogeneity, which we extracted using the `getCorrected()` function. These spatially-corrected plot-level values remove field position effects while preserving biological variation.

In the second stage, we modeled the spatially-corrected phenotypes 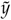 with a linear model to estimate treatment and genotype effects:

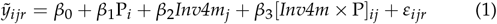

where the spatially-corrected observation 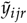 corresponds to plot *r* under phosphorus treatment *i* with genotype *j*. The fixed effects coefficients represent the overall mean (*β*_0_), the effect of phosphorus treatment (*β*_1_), the effect of genotype (*β*_2_), and their interaction (*β*_3_). The residuals 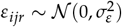 capture remaining variation after spatial correction. We adjusted p-values for multiple testing using the false discovery rate method and report effects with FDR *<* 0.05.

For each of the 16 field plant phenotype traits we extracted from Eq. 1 (`emmeans` (**?**)) the nine contrasts that span the 2 × 2 *Inv4m* × phosphorus design: the −P vs +P contrast at each genotype, the *Inv4m* vs CTRL contrast at each treatment, the two diagonal contrasts, the marginal Treatment and Genotype contrasts, and the Treatment × Genotype interaction. These 144 contrasts were combined into a single family prior to Benjamini Hochberg FDR adjustment. Figure 1 displays within-genotype contrasts for the traits with significant main effects; Supplementary Figure S1 Fig displays the marginal ±P contrasts for plant height, 50-kernel weight, total kernel number, and total kernel weight per plant.

### Tissue sampling, RNA extraction, and sequencing

We sampled the plants at 63 Days After Planting (DAP). We estimated the developmental stage to be around V13. Which meant that at the time of measurement, the plant had grown ~ 13 fully collared leaves, from which ~ 8 remained green. We sampled every other leaf downwards to maximize the developmental divergence between our allotted four samples per individual and represent the physiological age gradient of the available mature canopy. We took tissue from the collar leaf (the first leaf with a fully developed collar, designated as Node 0 in (Ciganda *et al*. 2008)), and continued sampling every other leaf below it, resulting in a total of four sampled leaves per plant. Specifically, the four sampled leaves corresponded to the following nodal positions: Node 0 (Collar Leaf), Node 2, Node 4, and Node 6 (with Node 6 being the most basal/oldest leaf sampled). These leaves were numbered sequentially from 1(most apical, Node 0) to 4 (most basal, Node 6) for analysis purposes. We used four replicate plants per combination of P treatment and *Inv4m* genotype for a total of 64 tissue samples. We took ten disc samples from the leaf tips with a tissue puncher and immediately froze the tissue in 1.5 mL tubes with two steel beads precooled with liquid nitrogen and kept in dry ice until stored at −80 °C. We extracted total RNA with the QIAGEN RNAeasy Plant Mini Kit RNA extraction kit following manufacturer procedures (QIAGEN 74904), and RNA samples were quantified in nanodrop and sent to the NCSU Core Genomics Laboratory for sequencing. Following QC in the Bioanalyzer, Illumina libraries were prepared and sequenced in a lane of the Novaseq according to the manufacturer’s recommendations.

### Lipid extraction, identification, and quantification by HPLC-MS

We used the lipid extraction by methyltert-butyl ether matyash2008 and HPLC-MS methods as described in (Barnes *et al*. 2022). First, we ground the frozen tissue samples using a SPEX Geno/Grinder (Metuchen, NJ, USA). Briefly, cold methanol (MeOH) and internal standard mix were added to the ground frozen tissue. This was vortexed and methyl tert-butyl ether (MTBE) was added before vortexing again. After shaking the samples at 4°C, we added LC/MS grade water at room temperature and vortexed. We then centrifuged the samples and collected the supernatant from the upper organic phase, splitting it into two aliquots and drying it in a rotovap. Dry samples were resuspended in 110 *µ*L of 100% MeOH (with the internal standard CUDA at 50 ng/mL), vortexed at low speed for 20 s, and then sonicated at room temperature for 5 min. We transferred the samples into amber glass vials with inserts prior to analysis. Lipid profiling was performed using a Thermo Scientific Orbitrap Exploris 480 mass spectrometer coupled to a Thermo Vanquish Horizon UHPLC. Mass spectrometry parameters and details were repeated as in (Barnes *et al*. 2022). We converted the RAW files to ABF and imported them into MS-DIAL version 4.24. The parameters were as follows: MS1 tolerance 0.01 Da, MS2 tolerance 0.025 Da, Minimum peak height 100,000, mass slice width 0.05 Da. A library of retention times from our standards was used for retention time identification, and the ID score cutoff was 80%. We also used the internal MS-DIAL compound library from the Fiehn laboratory with a retention time tolerance of 1 min. For alignment, all samples were compared to one of the pooled QCs. Quantification was calculated as the area under the peak. Any internal standard, non-plant lipid, or odd-chain fatty acid-containing lipid was removed from quantification.

### Differential gene expression and differential lipid analysis

We aligned RNA-seq reads to the maize Zm-B73-REFERENCE-NAM-5.0 genome using *kallisto* bray2016. Transcript-level alignments were aggregated to gene-level counts per biological replicate. Genes with low expression across samples were filtered using `filterByExpr` from *edgeR* robinson2010. For lipid analysis, we removed internal standards and odd-chain fatty acid species, then processed MS-Dial peak area data. Lipid ion counts were scaled by total ion count and multiplied by 1 × 10^9^ for numerical stability, then analyzed without further normalization.

We applied TMM normalization to gene expression counts to adjust for sequencing depth differences. For both gene expression and lipid abundance data, we calculated variance weights with *voom* law2014 to model the mean-variance relationship and transformed data to log_2_ scale (log_2_(CPM) for genes, log_2_(scaled abundance) for lipids). We fitted multivariate linear models separately for gene expression and lipid abundance using *limma* ritchie2015.

For the log-transformed response *Y*_*ijrs*_ (expression or abundance) from leaf stage *s*, in plant *r*, under phosphorus treatment *i*, with genotype *j*, we have:

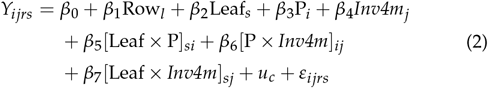

with random effect and residuals:

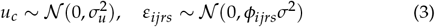

where Leaf_*s*_ represents the centered continuous leaf stage variable (*s* ∈ {1, 2, 3, 4}, centered at mean 2.5), such that *β*_2_ represents the rate of change per leaf stage increment. The term *u*_*c*_ is a random effect for plot column nested within treatment, accounting for spatial autocorrelation. The residual variance *ϕ*_*ijrs*_*σ*^2^ incorporates heteroscedastic precision weights from *voom*. Lipid models additionally included MS injection order as a technical covariate (S9 Fig). Empirical Bayes moderation was applied using `eBayes` to stabilize variance estimates across genes or lipids.

We adjusted p-values for multiple testing as false discovery rates (FDR). For phosphorus treatment and *Inv4m* genotype main effects, genes or lipids with FDR *<* 0.05 and | log_2_(FC)| *>* 1.5 were classified as strong DEGs or strong DALs. For leaf stage and interaction effects, we used a threshold of | log_2_(FC)| *>* 0.5, equivalent to | log_2_(FC)| *>* 1.5 between leaf stages 1 and 4. R scripts and expression data are available at the inv4m GitHub repository.

### Gene Ontology and KEGG overrepresentation analysis

Once we had sets of differentially expressed genes for the three predictors (leaf, −P, *Inv4m*) and two types of gene expression response (upregulated and downregulated), we proceeded to annotate them with gene ontology terms and KEGG pathways using *ClusterProfiler* (Yu *et al*. 2012; Zicola 2024). We started with the B73 NAM v5 gene ontology annotation from (Fattel *et al*. 2024) and added GO terms for each intermediate node in the gene ontology tree using the *ClusterProfiler* function *buildGOmap*. Then, we conducted gene over-representation analysis using the function *compareCluster*, with the set of 24011 genes detected in at least one high-quality leaf RNA-seq library as the universe/background. This function calculates the hypergeometric test for overrepresented ontology terms in the specified gene set and returns raw and FDR-adjusted p-values. We then manually reviewed the combined 1700 significant (*FDR <* 0.05) overrepresented GO term associations for the 6 predictor/regulation combinations. For illustration, we selected an *ad hoc* subset representative of biological relevance and minimizing semantic redundancy. Similarly, we tested for KEGG pathway overrepresentation using the *enrichKEGG* function from *compareCluster*, which performs the same hypothesis tests on the NCBI RefSeq annotation of the B73 NAM assembly. Both types of overrepresentation analysis were plotted with the package *DOSE* (Yu *et al*. 2015).

### Gene Set Transcriptomic Indices

To visualize developmental and stress-induced changes in major biological processes, we calculated transcriptomic indices for four gene sets. The Chlorophyll Synthesis set was compiled from CornCyc pathway annotations. The Chlorophyll Degradation set was manually curated from the same CornCyc reactions, supplemented with six additional genes drawn from KEGG and (Wei *et al*. 2025) that mediate canonical chlorophyll catabolism in maize but were missing from the CornCyc reaction set. The Photosynthesis (163 genes) and Leaf Senescence (55 genes) sets were taken from (Ojeda-Rivera *et al*. 2026), who curated pani-coid grass orthologs of canonical photosynthesis and senescence regulators.

For each gene set, we calculated the mean log_10_(CPM) across all genes in the set for each sample. Index values were then normalized to represent the proportion of the total expression range across the four leaf stages using the formula:

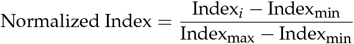

where Index_*i*_ represents the mean expression for leaf stage *i*, and Index_min_ and Index_max_ represent the minimum and maximum mean expression values across all four leaf stages. This normalization scales all indices to a 0-1 range, facilitating visual comparison of expression trajectories across gene sets with different baseline expression levels.

## Acknowledgments

Fieldwork and mapping population development were supported by NSF-PGR award 1546719 and USDA awards 2022-67013-37038 and 2022-67013-38264. This work is supported by the Research Capacity Fund (HATCH), project award nos. 7005660 and 7008935, from the U.S. Department of Agriculture’s National Institute of Food and Agriculture. The work on this paper and Nirwan Tandukar was supported by the U.S. Department of Energy, Office of Science, Biological and Environmental Research program, Early Career Award Number DE-SC0021889. Allison Barnes was supported by NSF-PGRP PRFB grant 2010703. Fausto Rodríguez-Zapata was supported by the Science and Technologies for Phosphorus Sustainability (STEPS) Center, a National Science Foundation Science and Technology Center (CBET-2019435). This work was performed in part by the Molecular Education, Technology and Research Innovation Center (METRIC) at NC State University, which is supported by the State of North Carolina. We thank members of the Ruairidh Sawers and Rubén Rellán Álvarez labs for their contribution to population development and field work that made possible the research in this manuscript. We thank the Puerto Vallarta Winter Nursery crews who have helped generate introgression populations used in this manuscript. We thank the staff at Penn State’s Rock Springs and NC State’s Central Crops Research Stations for supporting field work shown in this paper. We especially want to acknowledge the indigenous people of the Americas and the ingenuity with which they domesticated and facilitated the spread and adaptation of maize throughout the continent. This work would not have been possible without the international maize research community and the willingness of so many colleagues to support the development of new research programs. Any opinions, findings, conclusions, or recommendations expressed in this publication are those of the author(s) and should not be construed to represent any official USDA, NSF, DOE, ARS or U.S. Government determination or policy.

## Supplement

**S1 Fig.**
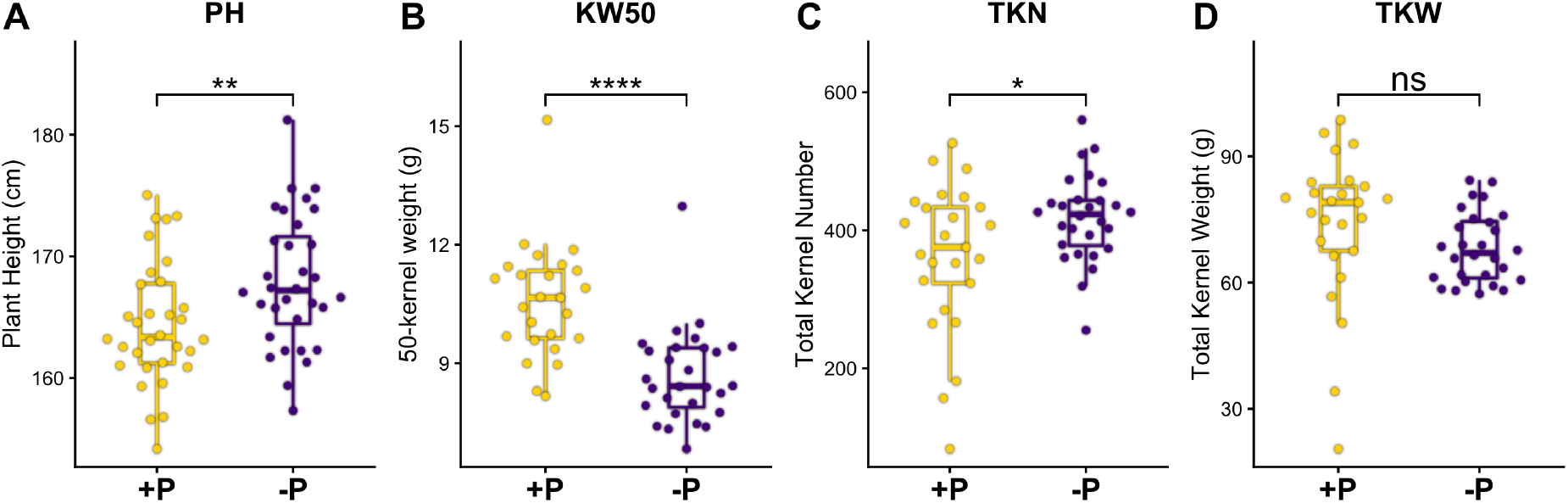
Marginal effects of phosphorus deficiency on plant height and kernel traits. Boxplots show individual field plot values for +P and −P treatments pooled across CTRL and *Inv4m* genotypes. **(A)** Plant height decreased under −P. **(B)** Fifty-kernel weight (a proxy for kernel size) decreased significantly under phosphorus deficiency. **(C)** Total kernel number showed a marginal increase under −P, consistent with a kernel size-to-number trade-off that collectively preserved total kernel weight. **(D)** Total kernel weight per plant was statistically unchanged under −P, indicating that kernel mass per plant was maintained at the expense of individual kernel size. *FDR* adjusted contrast *t-test* significance: *n*.*s*. not significant, *p <* 0.05 (*), *p <* 0.01 (**), *p <* 0.001 (***), *p <* 0.0001 (****).

**S2 Fig.**
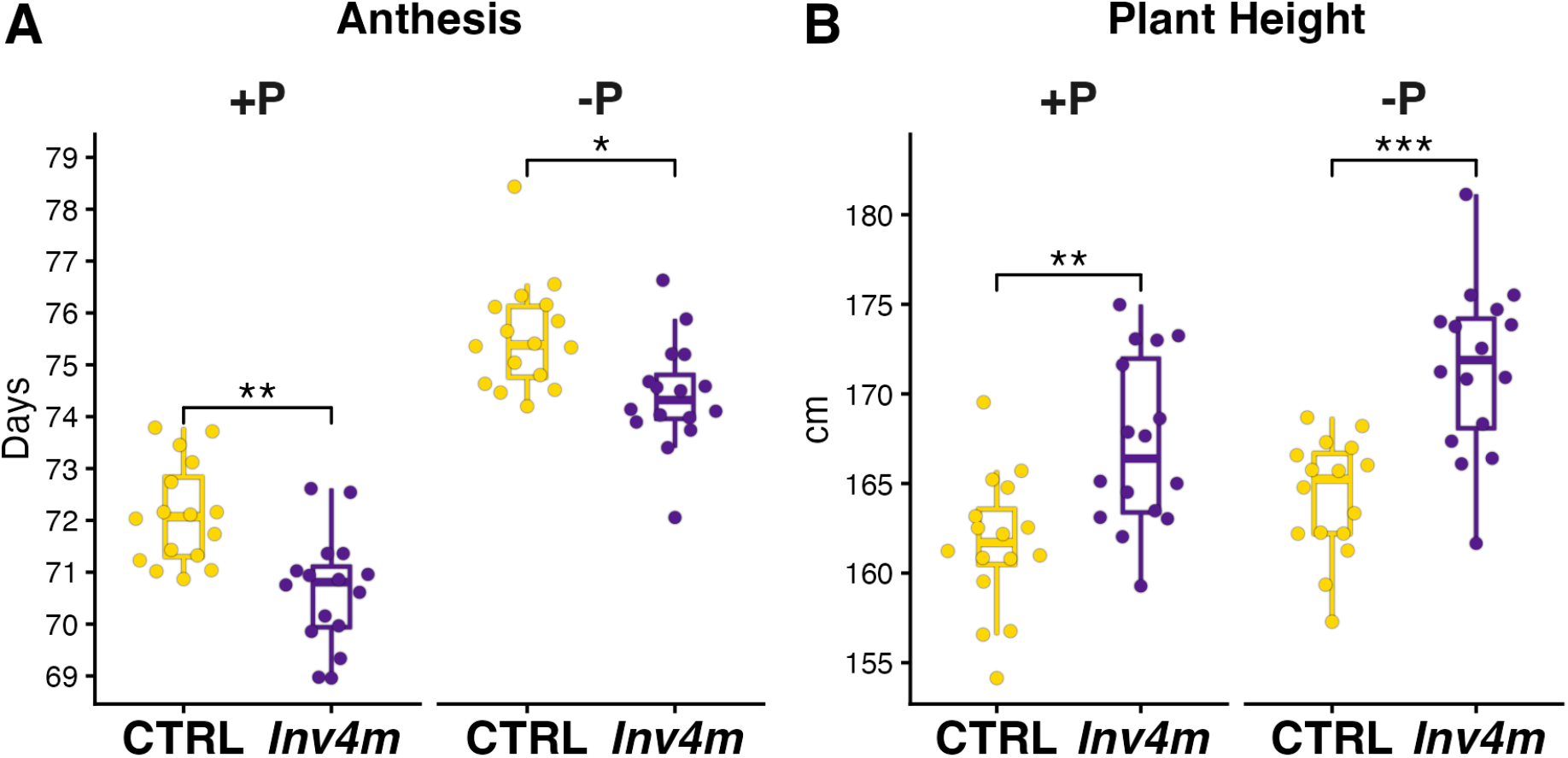
*Inv4m* differences in Anthesis and Plant Height. **(A)** Days to anthesis. *Inv4m* reached anthesis significantly earlier than CTRL under both phosphorus sufficient (+P) and deficient (−P) conditions. **(B)** Plant height at sampling. *Inv4m* plants were significantly taller than CTRL plants under both +P and −P treatments. Yellow boxplots represent CTRL, purple boxplots represent *Inv4m. FDR* adjusted contrast *t-test* significance: *n*.*s*. not significant, *p <* 0.05 (*), *p <* 0.01 (**), *p <* 0.001 (***), *p <* 0.0001 (****).

**S3 Fig.**
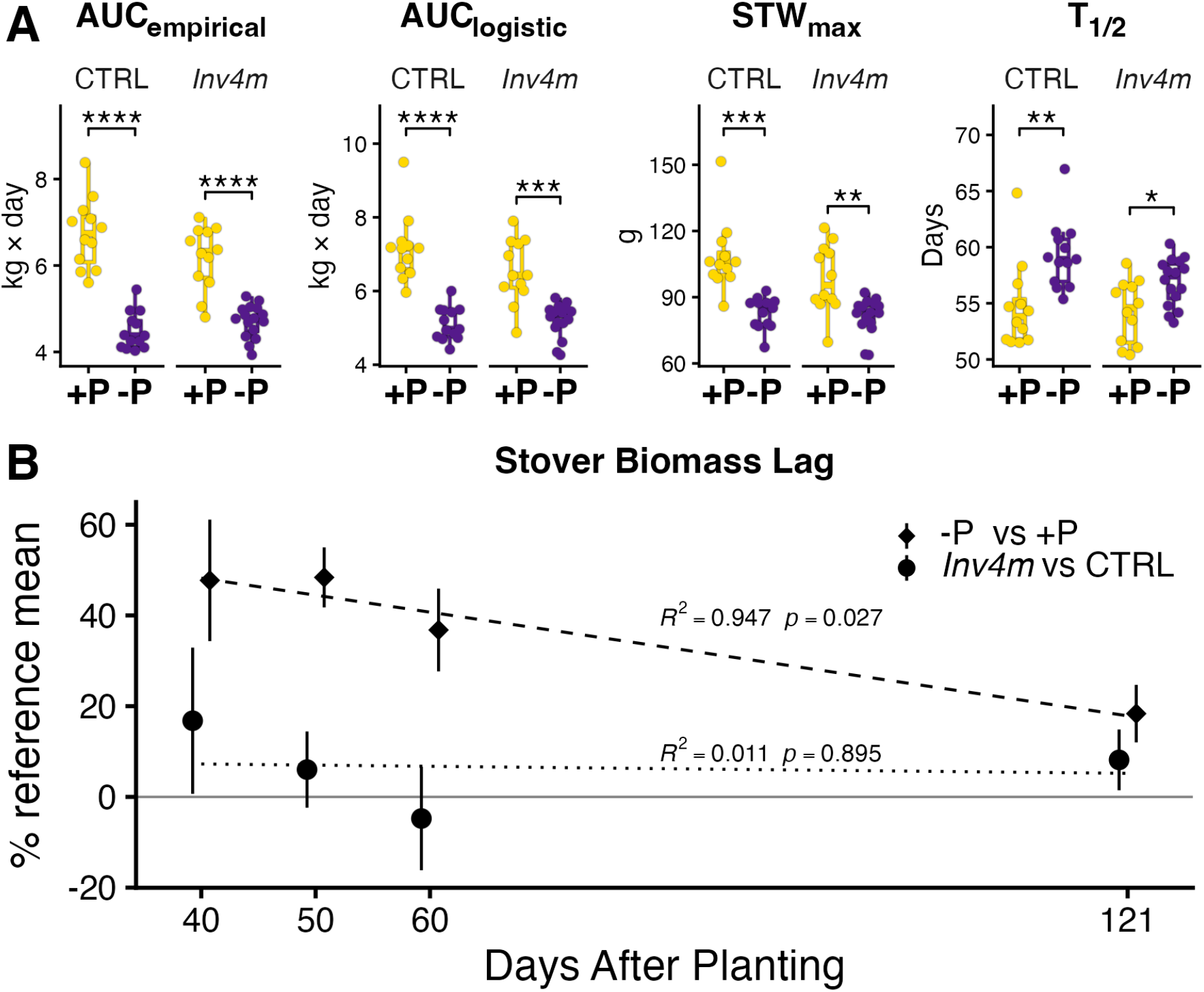
Maize Stover Dry Weight Growth Curves and Derived Parameters Highlight Phosphorus-Dependent Effects with No Genotype-by-Environment Interaction. **(A)** Derived logistic growth parameters for CTRL and *Inv4m* genotypes under +P and −P. Phosphorus deficiency significantly reduced the Area Under the Curve (AUC) for empirical data **(B)** and logistic model **(C)**, prolonged the time to reach half maximum stover weight (T_1/2_) (D), and decreased the maximum stover weight (STW_max_) (E). *FDR* adjusted Welch’s *t-test* significance: *n*.*s*. not significant, *p <* 0.05 (*), *p <* 0.01 (**), *p <* 0.001 (***), *p <* 0.0001 (****).

**S4 Fig.**
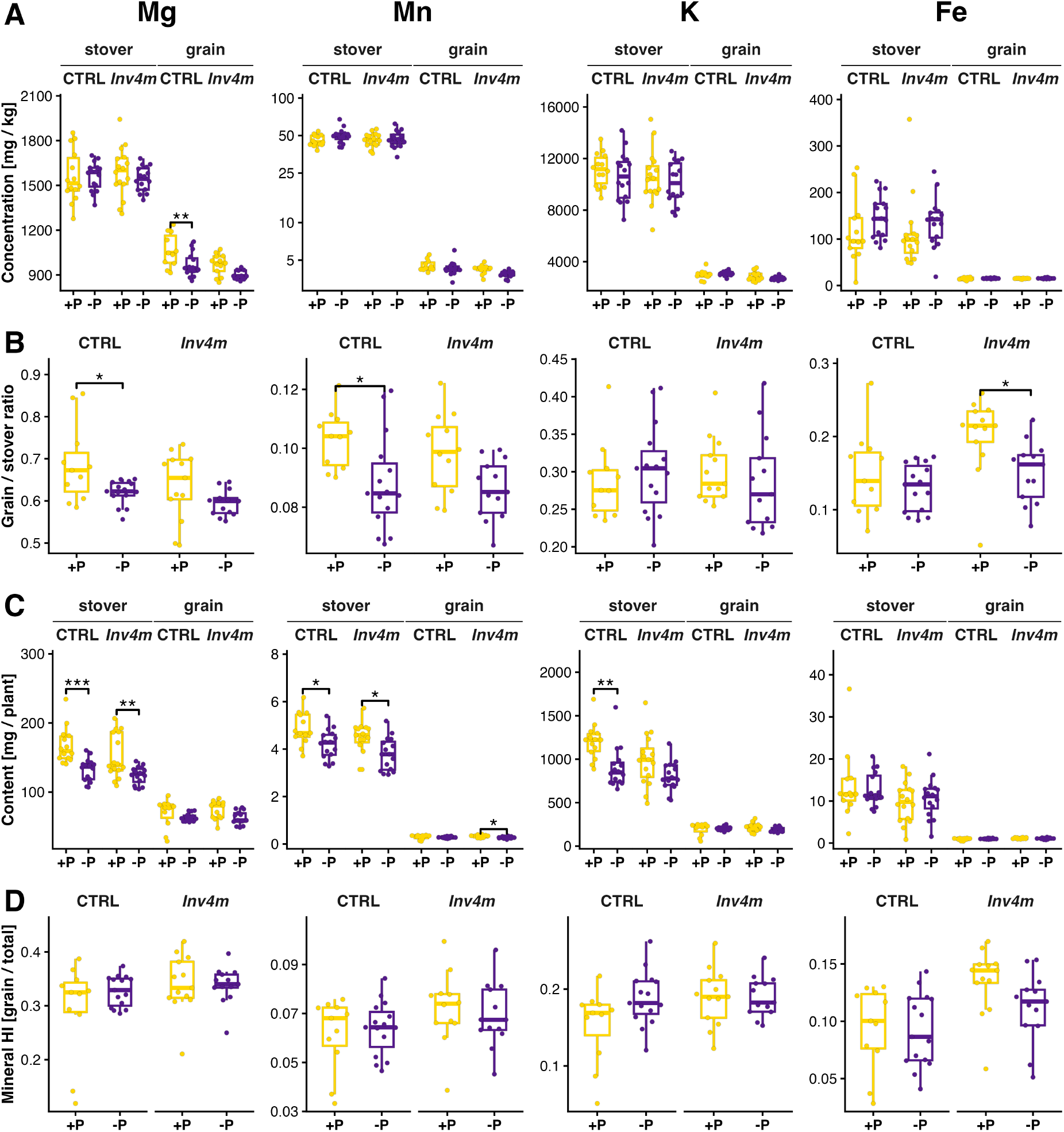
Secondary ionomic responses of *Inv4m* and control (CTRL) maize lines under phosphorus sufficiency (+P) and deficiency (−P). Boxplots for magnesium (Mg), manganese (Mn), potassium (K), and iron (Fe) show, per genotype, **(A)** element concentrations (mg/kg) in stover and grain, **(B)** grain/stover concentration ratios, **(C)** per plant content (mg/plant) in stover and grain, and **(D)** mineral harvest index (HI, grain/total). *FDR* adjusted contrast *t-test* significance: *n*.*s*. not significant, *p <* 0.05 (*), *p <* 0.01 (**), *p <* 0.001 (***), *p <* 0.0001 (****). Effect sizes and exact *p* values are reported in S1 Table.

**S5 Fig.**
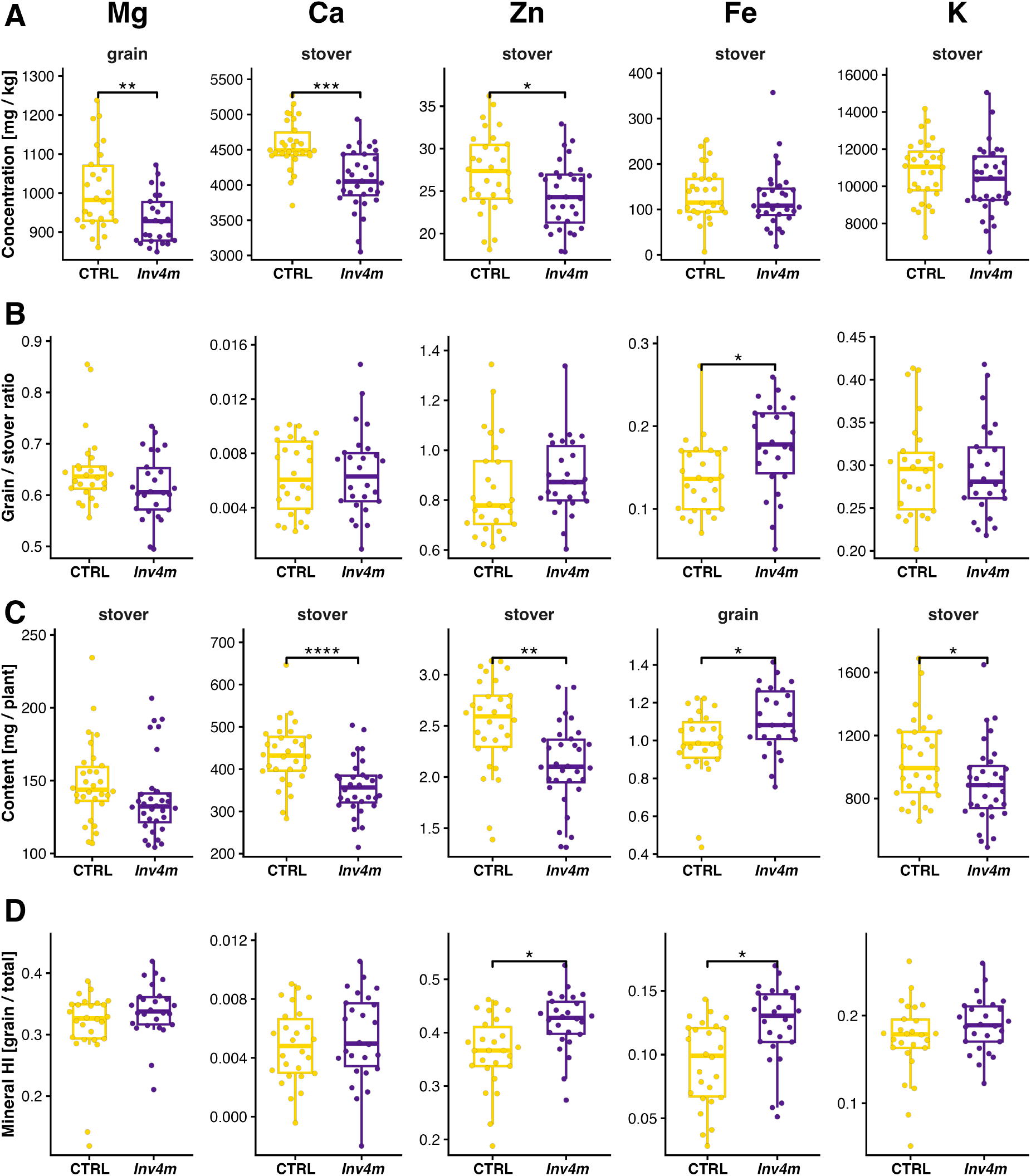
*Inv4m* effects on the ionome, pooled across phosphorus treatments. For each of five minerals (Mg, Ca, Zn, Fe, K; columns), boxplots contrast control (CTRL) and *Inv4m* plants on four metrics: **(A)** mineral concentration (mg/kg), **(B)** grain/stover concentration ratio, **(C)** per-plant mineral content (mg/plant), and **(D)** mineral harvest index (HI, grain/total). For concentration and content, a single pool is shown per cell: the pool where the *Inv4m* vs CTRL contrast is significant under the unified FDR, or *stover* when neither pool is significant. The selected pool is indicated on the facet strip. *FDR* adjusted contrast *t-test* significance: *n*.*s*. not significant, *p <* 0.05 (*), *p <* 0.01 (**), *p <* 0.001 (***), *p <* 0.0001 (****).

**S6 Fig.**
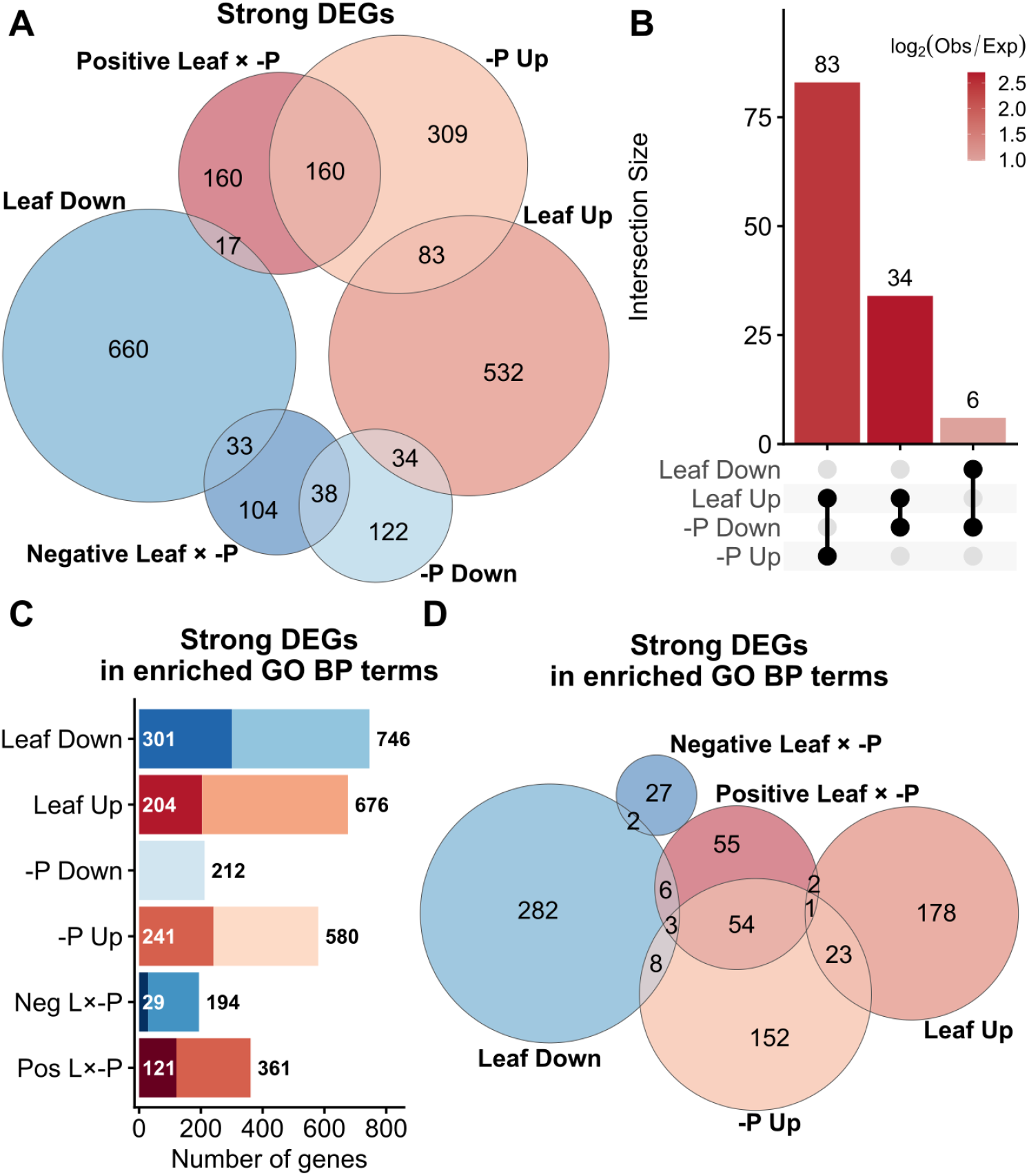
(**A**) Euler diagram showing the major two-way intersections among DEGs from the effects of leaf position (Leaf Up/Leaf Down), phosphorus deficiency (−P Up / −P Down), and interaction (Negative and Positive). Circle sizes represent set sizes, and numbers indicate the counts in each intersection. (**B**) Upset plot highlighting strong DEG intersections that are significantly enriched relative to expectation (FDR <0.05), the first and second intersections are depicted in A, the third (Leaf Down ∩ −P Down) is not, due to its small size. (**C**) Summary of strong DEGs and their GO Biological Process (BP) annotation. Black numbers indicate the total number of strong DEGs per group; white numbers denote the subset annotated with significantly enriched BP terms. Neg L×–P: negative Leaf × −P interaction effect, and Pos L×–P: positive interaction. (**D**) Euler diagram showing overlap among the strong BP annotated DEG sets, illustrating shared and unique functional responses across leaf position, phosphorus deficiency, and their interaction.

**S7 Fig.**
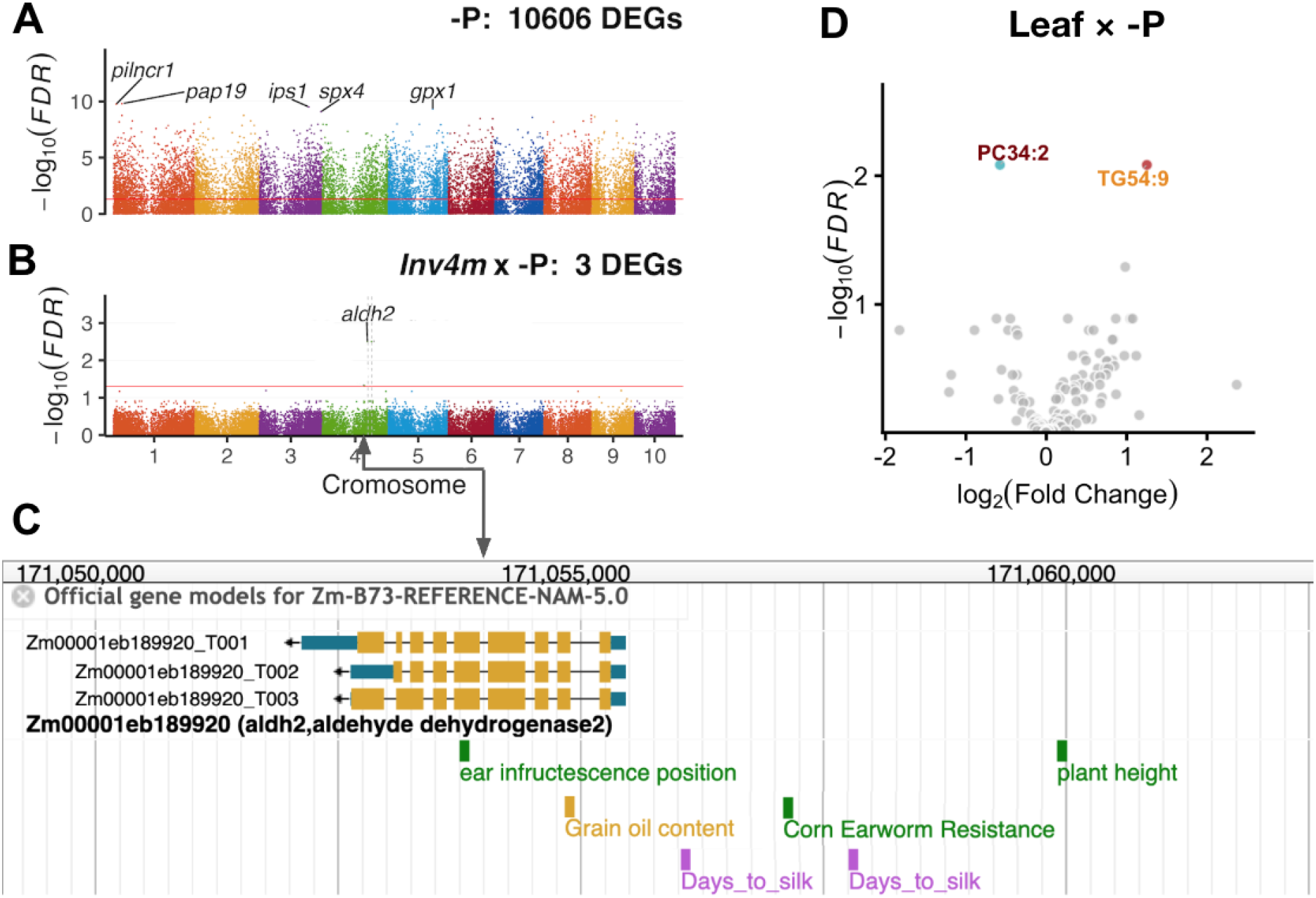
**(A) Manhattan plot for differentially expressed genes (DEGs) under −P.**Loci associated with key P-starvation response genes, such as *pilncr1, pap19, ips1*, and *spx4*, are highlighted. The *y*-axis represents the − log_10_(FDR). (**B**) **A single gene, *aldh2*, carries a significant *Inv4m*** × − **P interaction**. *aldh2* (*aldehyde dehydrogenase 2*) surpasses the *FDR <* 0.05 threshold and is located on Chromosome 4 outside the *Inv4m* inversion, within the flanking introgression. (**C**) Zoomed-in view of the *aldh2* locus on Chromosome 4 around position 171.05 Mb. (**D**) **Volcano plot for differentially abundant lipids (DALs) under Leaf** × − **P interaction**.

**S8 Fig.**
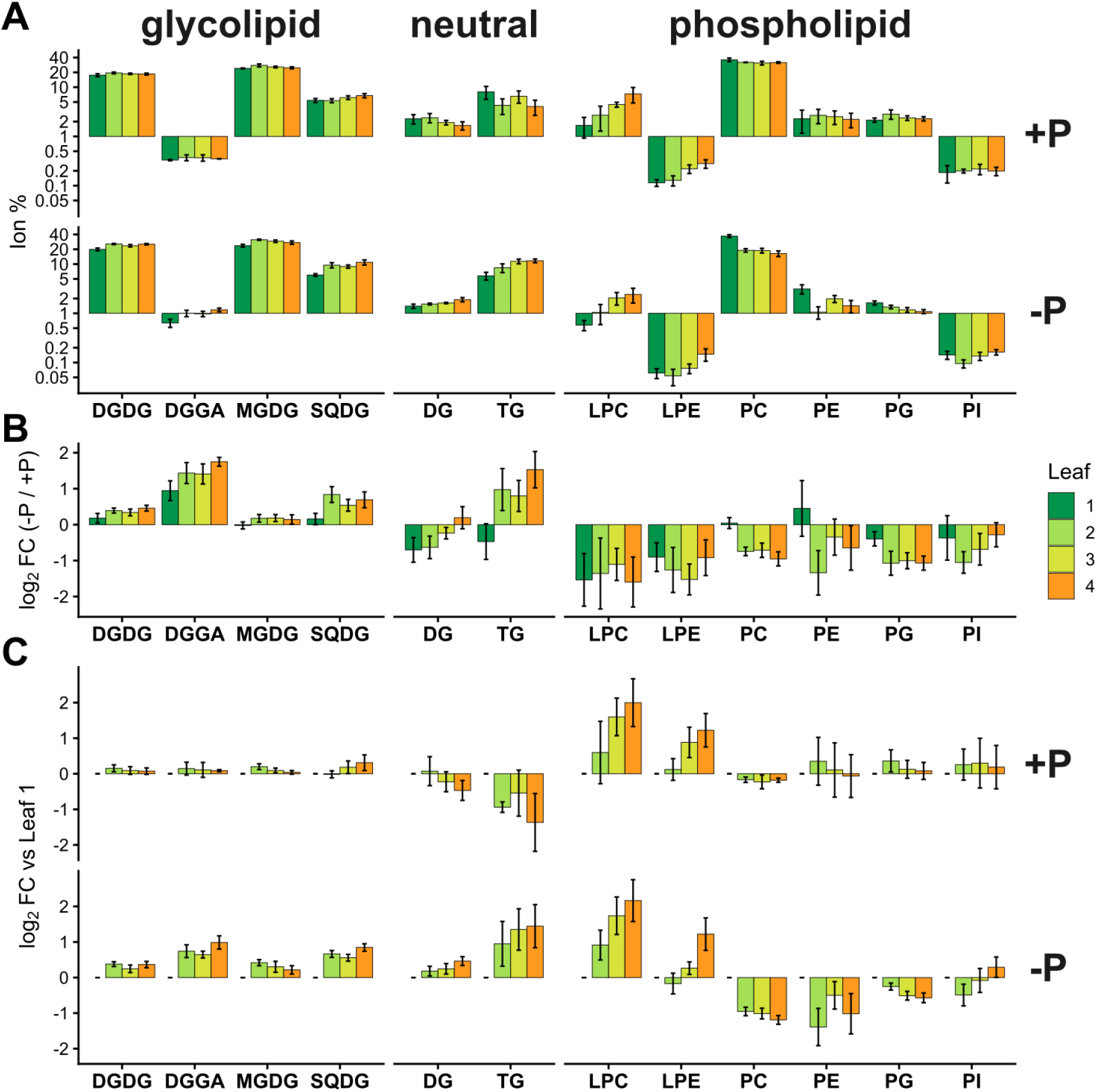
**Lipid class composition, and effect of leaf developmental stages and phosphorus treatments**. of **(A)** Membrane lipid composition (Ion %, log scale) is dominated by PC, MGDG, DGDG, and SQDG, while LPE, PI and DGGA (in +P) represent minor components (<1%, necessitating log scale). **(B)** Treatment effect (−P / +P) on lipids, as *log*_2_Fold Change, shows leaf-stage-dependent responses in DG, TG, PC, and PE. **(C)** leaf effect, as *log*_2_Fold Change, Error bars indicate standard error of the mean.

**S9 Fig.**
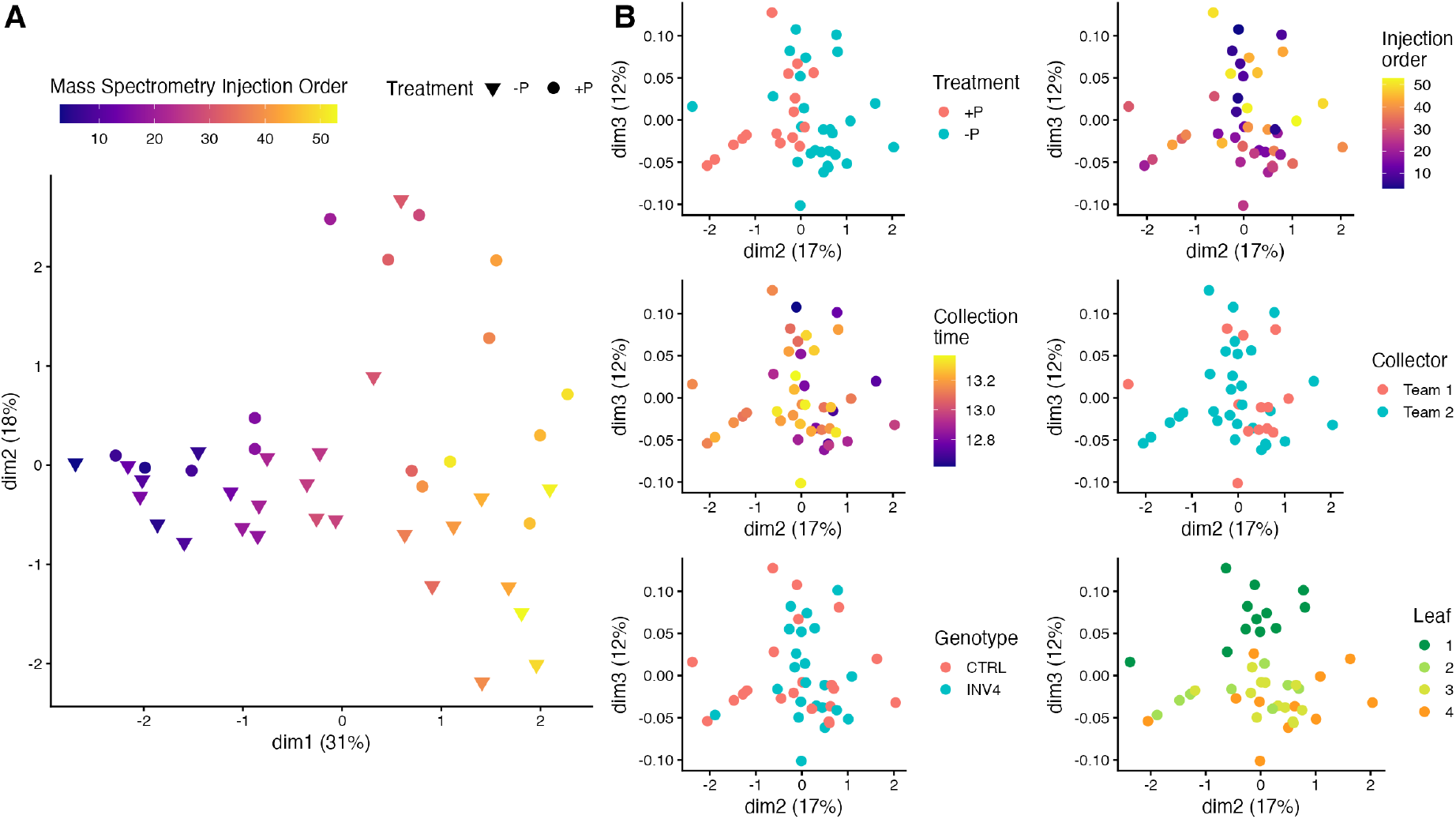
Mass spectrometry injection order confounds lipid profile variation. **(A)** Multidimensional scaling (MDS) of lipid profiles. Treatment groups (−P, +P) are distinguished by shape, while color gradient represents injection order (purple = early, yellow = late). The injection order was used as a covariate in the *limma* mixed linear model of lipid variation; see Methods. **(B)** Lipid data projections on the next most explanatory MDS dimensions (dim2 vs. dim3) colored by experimental factors.

**S1 Table.** Within-genotype −P vs +P ionome contrasts at *FDR<* 0.05. Significant rows from the 96-test `emmeans` family (48 mineral × metric responses × 2 within-genotype contrasts) underlying Figure 2 and Supplementary Figure S4 Fig. Estimates are −P vs +P marginal-mean differences in the native unit of each metric (ppm for concentration, mg/plant for content, dimensionless for grain/stover ratio and harvest index). Benjamini–Hochberg FDR across 96 tests.

| Metric | Mineral | Pool | Genotype | Estimate ± SE | FDR |
| --- | --- | --- | --- | --- | --- |
| Concentration | P | grain | CTRL | $-672 \pm 94$ | $3.9 \times 10^{-8}$ |
| | P | grain | *Inv4m* | $-684 \pm 93$ | $2.0 \times 10^{-8}$ |
| | P | stover | CTRL | $-1592 \pm 85$ | $1.1 \times 10^{-24}$ |
| | P | stover | *Inv4m* | $-1673 \pm 82$ | $3.1 \times 10^{-26}$ |
| | Zn | stover | CTRL | $6.85 \pm 1.12$ | $7.7 \times 10^{-7}$ |
| | Zn | stover | *Inv4m* | $5.02 \pm 1.08$ | $1.0 \times 10^{-4}$ |
| | Ca | grain | CTRL | $19.0 \pm 3.4$ | $9.6 \times 10^{-6}$ |
| | Ca | grain | *Inv4m* | $16.7 \pm 3.4$ | $6.4 \times 10^{-5}$ |
| | S | grain | CTRL | $79.1 \pm 29.9$ | 0.028 |
| | S | stover | CTRL | $113 \pm 21$ | $9.6 \times 10^{-6}$ |
| | Mg | grain | CTRL | $-97.5 \pm 30.5$ | 0.008 |
| Grain/stover ratio | P | – | CTRL | $2.09 \pm 0.16$ | $1.1 \times 10^{-15}$ |
| | P | – | *Inv4m* | $2.00 \pm 0.16$ | $2.2 \times 10^{-15}$ |
| | Zn | – | CTRL | $-0.306 \pm 0.049$ | $1.1 \times 10^{-6}$ |
| | Zn | – | *Inv4m* | $-0.178 \pm 0.049$ | 0.003 |
| | Ca | – | CTRL | $0.00416 \pm 0.00087$ | $9.2 \times 10^{-5}$ |
| | Ca | – | *Inv4m* | $0.00366 \pm 0.00086$ | $4.0 \times 10^{-4}$ |
| | Mg | – | CTRL | $-0.0673 \pm 0.0247$ | 0.025 |
| | Mn | – | CTRL | $-0.0149 \pm 0.0051$ | 0.017 |
| | Fe | – | *Inv4m* | $-0.0507 \pm 0.0180$ | 0.020 |
| Content | P | grain | CTRL | $-41.1 \pm 14.6$ | 0.020 |
| | P | grain | *Inv4m* | $-67.5 \pm 14.4$ | $1.1 \times 10^{-4}$ |
| | P | stover | CTRL | $-186 \pm 16$ | $1.1 \times 10^{-15}$ |
| | P | stover | *Inv4m* | $-169 \pm 15$ | $7.6 \times 10^{-15}$ |
| | Ca | grain | CTRL | $1.26 \pm 0.29$ | $3.4 \times 10^{-4}$ |
| | Ca | grain | *Inv4m* | $1.19 \pm 0.28$ | $4.8 \times 10^{-4}$ |
| | Ca | stover | CTRL | $-97.4 \pm 20.6$ | $8.0 \times 10^{-5}$ |
| | Ca | stover | *Inv4m* | $-67.1 \pm 19.9$ | 0.004 |
| | Mg | stover | CTRL | $-34.8 \pm 8.4$ | $4.2 \times 10^{-4}$ |
| | Mg | stover | *Inv4m* | $-27.5 \pm 8.1$ | 0.004 |
| | Mn | grain | *Inv4m* | $-0.061 \pm 0.023$ | 0.026 |
| | Mn | stover | CTRL | $-0.662 \pm 0.249$ | 0.026 |
| | Mn | stover | *Inv4m* | $-0.676 \pm 0.239$ | 0.020 |
| | K | stover | CTRL | $-299 \pm 84$ | 0.003 |
| Harvest index | P | – | CTRL | $0.280 \pm 0.031$ | $5.6 \times 10^{-11}$ |
| | P | – | *Inv4m* | $0.238 \pm 0.030$ | $3.9 \times 10^{-9}$ |
| | Ca | – | CTRL | $0.00373 \pm 0.00077$ | $8.0 \times 10^{-5}$ |
| | Ca | – | *Inv4m* | $0.00410 \pm 0.00076$ | $1.5 \times 10^{-5}$ |

**S2 Table.**
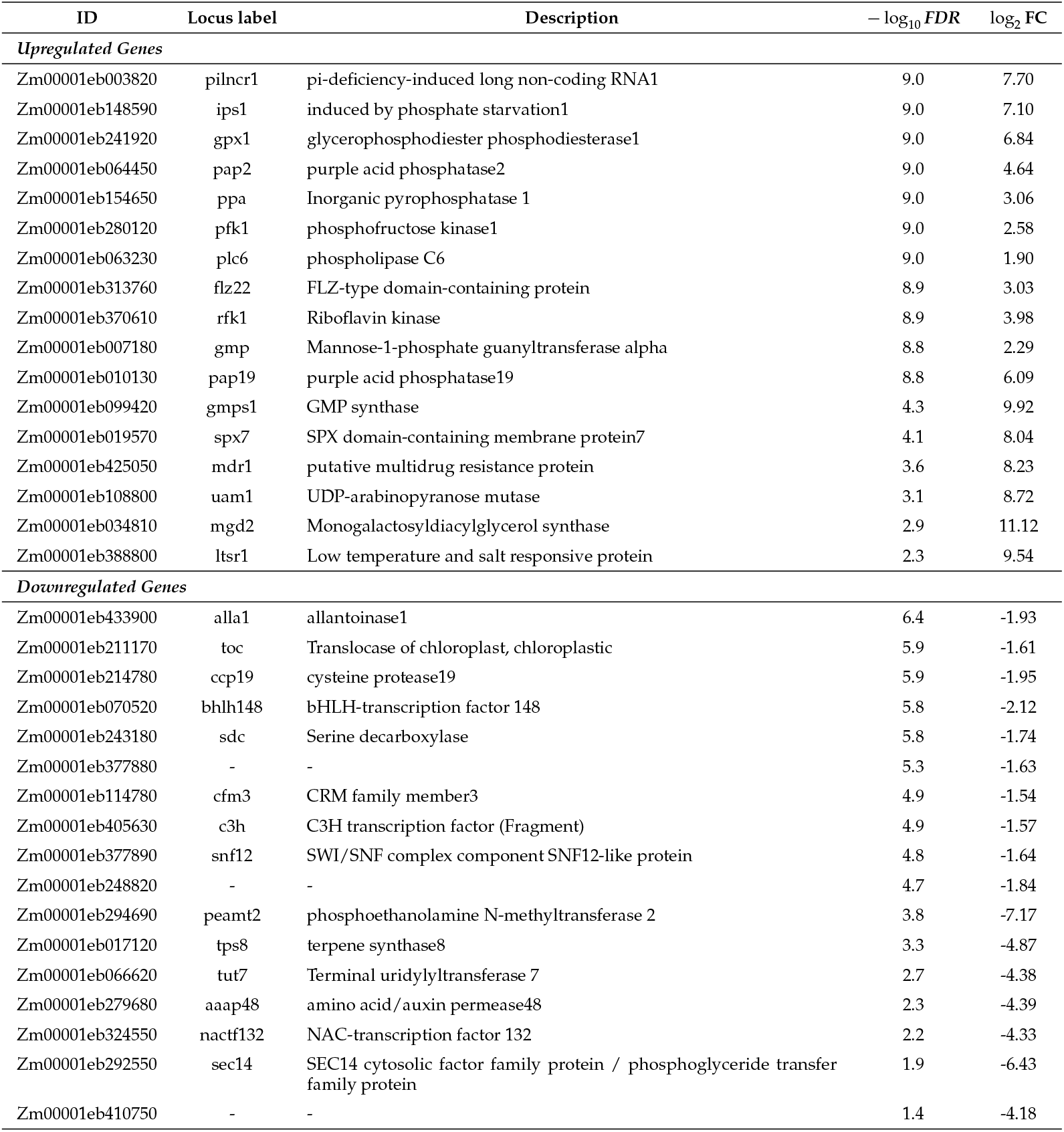
Selected Differentially Expressed Genes under Phosphorus Starvation (−P) effect.

**S3 Table.**
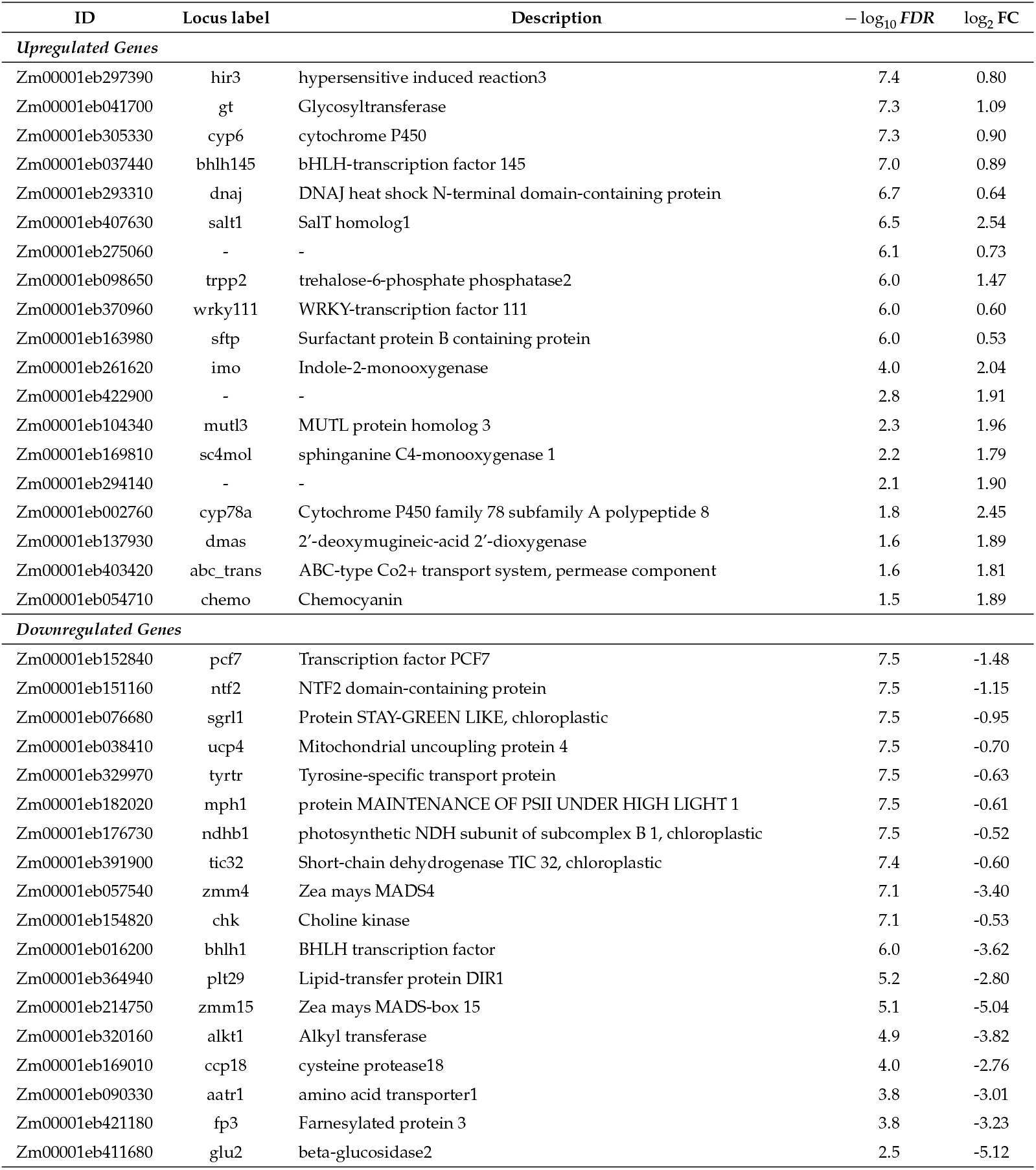
Selected Differentially Expressed Genes for Leaf Stage effect.

**S4 Table.** Selected Differentially Expressed Genes in Leaf × −P interaction, effect per increased Leaf Stage(−P).

| ID | Locus label | Description | $-\log_{10} FDR$ | $\log_2 FC$ |
| --- | --- | --- | --- | --- |
| *Positively Interacting Genes* |  |  |  |  |
| Zm00001eb157810 | pk | Pyruvate kinase | 5.6 | 1.18 |
| Zm00001eb376160 | mrpa3 | multidrug resistance-associated protein3 | 5.4 | 0.64 |
| Zm00001eb063230 | plc6 | phospholipase C6 | 4.5 | 0.56 |
| Zm00001eb144680 | rns | Ribonuclease T(2) | 4.4 | 0.61 |
| Zm00001eb339870 | pld16 | phospholipase D16 | 4.3 | 0.56 |
| Zm00001eb393060 | piplc | PI-PLC X domain-containing protein | 4.0 | 1.15 |
| Zm00001eb148030 | gmp1 | mannose-1-phosphate guanylyltransferase1 | 3.9 | 0.69 |
| Zm00001eb009430 | htm4 | Heptahelical transmembrane protein 4 | 3.9 | 0.63 |
| Zm00001eb011050 | bgal | Beta-galactosidase | 3.9 | 0.53 |
| Zm00001eb289800 | pah1 | phosphatidate phosphatase 1 | 3.9 | 0.58 |
| Zm00001eb263160 | ring | Zinc finger (C3HC4-type RING finger) family protein | 2.9 | 2.16 |
| *Negatively Interacting Genes* |  |  |  |  |
| Zm00001eb359280 | tat | Tat pathway signal sequence family protein | 5.6 | -0.56 |
| Zm00001eb207130 | cab | Chlorophyll a-b binding protein, chloroplastic | 5.4 | -1.35 |
| Zm00001eb389720 | fbpase | D-fructose-1,6-bisphosphate 1-phosphohydrolase | 5.3 | -0.81 |
| Zm00001eb070520 | bhlh148 | bHLH-transcription factor 148 | 5.1 | -0.96 |
| Zm00001eb212520 | psad1 | photosystem I subunit d1 | 4.6 | -0.62 |
| Zm00001eb179680 | cab | Chlorophyll a-b binding protein, chloroplastic | 4.6 | -0.55 |
| Zm00001eb111630 | med33a | Mediator of RNA polymerase II transcription subunit 33A | 4.4 | -0.60 |
| Zm00001eb362560 | ndho1 | NADH-plastoquinone oxidoreductase1 | 4.4 | -0.58 |
| Zm00001eb214780 | ccp19 | cysteine protease19 | 4.2 | -0.82 |
| Zm00001eb071770 | mex1 | maltose excess protein1 | 4.0 | -0.59 |
| Zm00001eb256120 |  |  | 3.8 | -1.41 |
| Zm00001eb235450 | taf2n | TATA-binding protein-associated factor 2N | 3.6 | -2.07 |
| Zm00001eb138960 |  |  | 2.1 | -2.11 |

**S5 Table.** Strong −P upregulated DEGs annotated with GO:0016036 “cellular response to phosphate starvation” ((Fattel *et al*. 2024))

| ID | Locus label | Description | $-\log_{10} FDR$ | $\log_2 FC$ |
| --- | --- | --- | --- | --- |
| *Upregulated Genes* |  |  |  |  |
| Zm00001eb241920 | gpx1 | glycerophosphodiester phosphodiesterase1 | 8.96 | 6.84 |
| Zm00001eb154650 | ppa1 | Inorganic pyrophosphatase 1 | 8.96 | 3.06 |
| Zm00001eb280120 | pfk1 | phosphofructose kinase1 | 8.96 | 2.58 |
| Zm00001eb370610 | rfk1 | Riboflavin kinase | 8.88 | 3.98 |
| Zm00001eb297970 | sqd2 | Sulfoquinovosyl transferase SQD2 | 8.05 | 1.83 |
| Zm00001eb347070 | sqd1 | sulfolipid biosynthesis1 | 8.05 | 1.76 |
| Zm00001eb335670 | sqd3 | Sulfoquinovosyl transferase SQD2 | 7.95 | 4.17 |
| Zm00001eb144680 | rns1 | Ribonuclease T(2) | 7.95 | 1.87 |
| Zm00001eb162710 | spx4 | SPX domain-containing membrane protein4 | 7.87 | 5.26 |
| Zm00001eb386270 | spx6 | SPX domain-containing membrane protein6 | 7.72 | 4.71 |
| Zm00001eb036910 | gpx3 | glycerophosphodiester phosphodiesterase3 | 7.72 | 3.50 |
| Zm00001eb151650 | pap1 | purple acid phosphatase1 | 7.37 | 4.61 |
| Zm00001eb351780 | ugp3 | UTP-glucose-1-phosphate uridylyltransferase 3, chloroplastic | 7.10 | 2.12 |
| Zm00001eb361620 | ppa2 | Inorganic pyrophosphatase 1 | 6.80 | 3.80 |
| Zm00001eb222510 | pht1 | phosphate transporter protein1 | 6.77 | 2.29 |
| Zm00001eb247580 | ppck3 | phosphoenolpyruvate carboxylase kinase3 | 6.34 | 2.96 |
| Zm00001eb126380 | phos1 | phosphate transporter1 | 6.12 | 2.17 |
| Zm00001eb038730 | pht7 | phosphate transporter protein7 | 5.68 | 6.56 |
| Zm00001eb116580 | spd1 | Protein seedling plastid development 1 | 5.60 | 1.66 |
| Zm00001eb048730 | spx2 | SPX domain-containing membrane protein2 | 5.42 | 4.90 |
| Zm00001eb069630 | oct4 | Organic cation/carnitine transporter 4 | 4.90 | 1.76 |
| Zm00001eb083520 | dgd1 | Digalactosyldiacylglycerol synthase | 4.85 | 1.77 |
| Zm00001eb430590 | nrx3 | Putative nucleoredoxin 3 | 4.71 | 4.15 |
| Zm00001eb130570 | sag21 | Senescence-associated gene 21, mitochondrial | 4.43 | 1.94 |
| Zm00001eb258130 | mgd3 | Monogalactosyldiacylglycerol synthase | 4.37 | 1.61 |
| Zm00001eb406610 | glk4 | G2-like-transcription factor 4 | 4.31 | 5.67 |
| Zm00001eb239700 | ppa2 | Inorganic pyrophosphatase 2 | 3.20 | 2.67 |
| Zm00001eb034810 | mgd2 | Monogalactosyldiacylglycerol synthase | 2.90 | 11.12 |
| Zm00001eb144670 | rns2 | Ribonuclease T(2) | 2.73 | 1.54 |
| Zm00001eb087720 | pht13 | phosphate transporter protein13 | 2.36 | 1.75 |
| Zm00001eb047070 | pht2 | phosphate transporter protein2 | 2.33 | 4.31 |
| Zm00001eb277280 | gst19 | glutathione transferase19 | 2.05 | 1.66 |
| Zm00001eb041390 | rns3 | Ribonuclease T(2) | 1.80 | 4.15 |
| Zm00001eb202100 | pap14 | purple acid phosphatase14 | 1.54 | 2.82 |

**S6 Table.** Strong DEGs with positive Leaf × −P interaction annotated with GO:0016036 “cellular response to phosphate starvation” ((Fattel *et al*. 2024))

| ID | Locus label | Description | $-\log_{10} FDR$ | $\log_2 FC$ |
| --- | --- | --- | --- | --- |
| *Upregulated Genes* |  |  |  |  |
| Zm00001eb144680 | rns1 | Ribonuclease T(2) | 4.39 | 0.61 |
| Zm00001eb339870 | pld16 | phospholipase D16 | 4.30 | 0.56 |
| Zm00001eb289800 | pah1 | phosphatidate phosphatase 1 | 3.86 | 0.58 |
| Zm00001eb297970 | sqd2 | Sulfoquinovosyl transferase SQD2 | 3.86 | 0.53 |
| Zm00001eb154650 | ppa1 | Inorganic pyrophosphatase 1 | 3.42 | 0.73 |
| Zm00001eb258130 | mgd3 | Monogalactosyldiacylglycerol synthase | 2.99 | 0.65 |
| Zm00001eb280120 | pfk1 | phosphofructose kinase1 | 2.99 | 0.56 |
| Zm00001eb335670 | sqd3 | Sulfoquinovosyl transferase SQD2 | 2.81 | 0.96 |
| Zm00001eb130570 | sag21 | Senescence-associated gene 21, mitochondrial | 2.75 | 0.73 |
| Zm00001eb430590 | nrx3 | Putative nucleoredoxin 3 | 2.46 | 1.40 |
| Zm00001eb151650 | pap1 | purple acid phosphatase1 | 2.45 | 1.09 |
| Zm00001eb369590 | nrx1 | Thioredoxin, nucleoredoxin | 2.21 | 0.83 |
| Zm00001eb238670 | pep2 | phosphoenolpyruvate carboxylase2 | 2.16 | 0.58 |
| Zm00001eb370610 | rfk1 | Riboflavin kinase | 2.01 | 0.66 |
| Zm00001eb247580 | ppck3 | phosphoenolpyruvate carboxylase kinase3 | 1.97 | 0.67 |
| Zm00001eb386270 | spx6 | SPX domain-containing membrane protein6 | 1.83 | 0.82 |
| Zm00001eb239700 | ppa2 | Inorganic pyrophosphatase 2 | 1.76 | 0.89 |
| Zm00001eb277280 | gst19 | glutathione transferase19 | 1.71 | 0.80 |
| Zm00001eb048730 | spx2 | SPX domain-containing membrane protein2 | 1.42 | 1.04 |
| Zm00001eb361620 | ppa2 | Inorganic pyrophosphatase 1 | 1.31 | 0.62 |

**S7 Table.** Strong DALs for leaf main effect.

| Lipid (IUB) | Class | $-\log_{10}(FDR)$ | $\log_2(FC)$ |
| --- | --- | --- | --- |
| *Accumulated Lipids* |  |  |  |
| LPC18:3 | phospholipid | 3.0 | 1.43 |
| LPE18:3 | phospholipid | 2.5 | 1.26 |
| PC36:6 | phospholipid | 2.5 | 0.73 |
| DGGA36:3 | glycolipid | 1.6 | 0.65 |
| LPE18:2 | phospholipid | 1.6 | 0.55 |
| PC38:5 | phospholipid | 1.6 | 0.67 |
| *Depleted Lipids* |  |  |  |
| DG36:4 | neutral | 4.9 | -0.71 |
| LPC16:0 | phospholipid | 2.9 | -1.36 |
| PE32:1 | phospholipid | 2.9 | -0.74 |
| DG34:2 | neutral | 2.5 | -0.51 |
| PE34:1 | phospholipid | 2.4 | -0.73 |
| PC36:4 | phospholipid | 2.2 | -0.50 |
| DGDG34:1 | glycolipid | 2.1 | -0.56 |
| DG26:0 | neutral | 2.0 | -0.68 |
| DG40:8 | neutral | 1.9 | -0.74 |
| PC36:1 | phospholipid | 1.8 | -0.69 |
| DGGA36:4 | glycolipid | 1.7 | -0.72 |
| DGDG36:1 | glycolipid | 1.6 | -4.09 |

**S8 Table.**
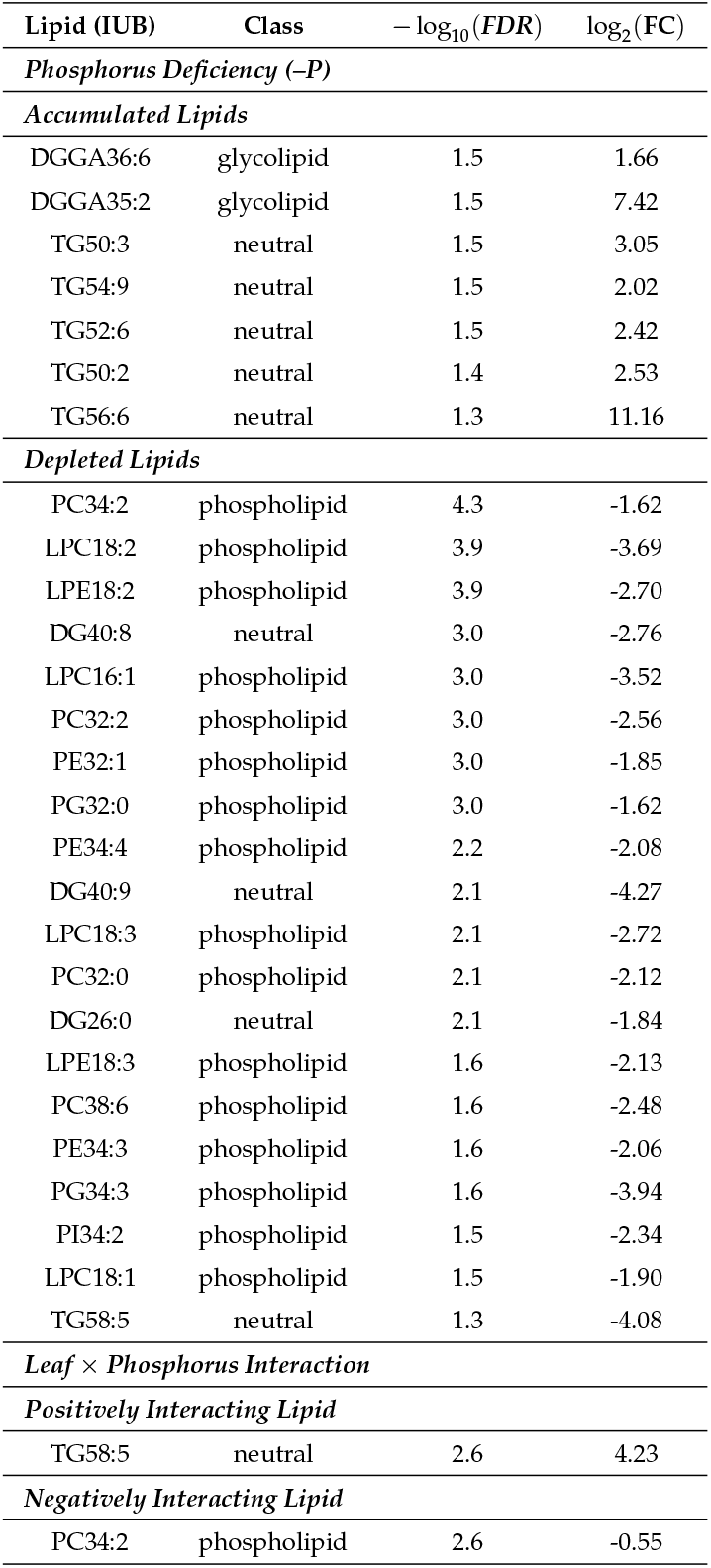
Strong DALs under phosphorus deficiency and its interaction with leaf stage.

| Lipid (IUB) | Class | $-\log_{10}(FDR)$ | $\log_2(FC)$ |
| --- | --- | --- | --- |
| *Phosphorus Deficiency (-P)* |  |  |  |
| *Accumulated Lipids* |  |  |  |
| DGGA36:6 | glycolipid | 1.5 | 1.66 |
| DGGA35:2 | glycolipid | 1.5 | 7.42 |
| TG50:3 | neutral | 1.5 | 3.05 |
| TG54:9 | neutral | 1.5 | 2.02 |
| TG52:6 | neutral | 1.5 | 2.42 |
| TG50:2 | neutral | 1.4 | 2.53 |
| TG56:6 | neutral | 1.3 | 11.16 |
| *Depleted Lipids* |  |  |  |
| PC34:2 | phospholipid | 4.3 | -1.62 |
| LPC18:2 | phospholipid | 3.9 | -3.69 |
| LPE18:2 | phospholipid | 3.9 | -2.70 |
| DG40:8 | neutral | 3.0 | -2.76 |
| LPC16:1 | phospholipid | 3.0 | -3.52 |
| PC32:2 | phospholipid | 3.0 | -2.56 |
| PE32:1 | phospholipid | 3.0 | -1.85 |
| PG32:0 | phospholipid | 3.0 | -1.62 |
| PE34:4 | phospholipid | 2.2 | -2.08 |
| DG40:9 | neutral | 2.1 | -4.27 |
| LPC18:3 | phospholipid | 2.1 | -2.72 |
| PC32:0 | phospholipid | 2.1 | -2.12 |
| DG26:0 | neutral | 2.1 | -1.84 |
| LPE18:3 | phospholipid | 1.6 | -2.13 |
| PC38:6 | phospholipid | 1.6 | -2.48 |
| PE34:3 | phospholipid | 1.6 | -2.06 |
| PG34:3 | phospholipid | 1.6 | -3.94 |
| PI34:2 | phospholipid | 1.5 | -2.34 |
| LPC18:1 | phospholipid | 1.5 | -1.90 |
| TG58:5 | neutral | 1.3 | -4.08 |
| *Leaf × Phosphorus Interaction* |  |  |  |
| *Positively Interacting Lipid* |  |  |  |
| TG58:5 | neutral | 2.6 | 4.23 |
| *Negatively Interacting Lipid* |  |  |  |
| PC34:2 | phospholipid | 2.6 | -0.55 |

**S1 File. Strong DEGs Associated with Senescence**. Strong DEGs that have been reported to be associated with senescence; they might respond to any of the experimental predictors in this study: −P, Leaf, *Inv4m* genotype.

## Notes

### Competing Interest Statement

The authors have declared no competing interest.

### Summary of Updates

In this revision we are simply updating a reference to include a link to a companion paper that we just uploaded to biorxiv: https://www.biorxiv.org/content/10.64898/2026.06.16.732082v1

